# Cellular senescence promotes progenitor cell expansion during axolotl limb regeneration

**DOI:** 10.1101/2022.09.01.506196

**Authors:** Qinghao Yu, Hannah E. Walters, Giovanni Pasquini, Sumeet Pal Singh, Martina Lachnit, Catarina Oliveira, Daniel León-Periñán, Andreas Petzold, Preethi Kesavan, Cristina Subiran, Ines Garteizgogeascoa, Dunja Knapp, Anne Wagner, Andrea Bernardos, María Alfonso, Gayathri Nadar, Alwin M. Graf, Konstantin E. Troyanovskiy, Andreas Dahl, Volker Busskamp, Ramón Martínez-Máñez, Maximina H. Yun

**Affiliations:** Technische Universita□t Dresden, CRTD/Center for Regenerative Therapies Dresden, Dresden, Germany; Department of Ophthalmology, Medical Faculty, University of Bonn, Bonn, Germany; IRIBHM, Université Libre de Bruxelles (ULB), Brussels, Belgium; Technische Universita□t Dresden, Center for Information Services and High Performance Computing (ZIH), Technische Universität Dresden, Dresden, Germany; DRESDEN-concept Genome Center (DcGC), Center for Molecular and Cellular Bioengineering, TU Dresden, Dresden, Germany; Instituto Interuniversitario de Investigación de Reconocimiento Molecular y Desarrollo Tecnológico (IDM), Universitat Politècnica de València, Valencia, Spain; CIBER de Bioingeniería, Biomateriales y Nanomedicina (CIBER-BBN), Instituto de Salud Carlos III; Unidad Mixta de Investigación en Nanomedicina y Sensores, Universitat Politècnica de València, IIS La Fe, Valencia, Spain; Max Planck Institute of Molecular Cellular Biology and Genetics, Dresden, Germany; Unidad Mixta UPV-CIPF de Investigación en Mecanismos de Enfermedades y Nanomedicina, Universitat Politècnica de València, Centro de Investigación Príncipe Felipe, Valencia, Spain; Physics of Life Excellence Cluster, Dresden, Germany

## Abstract

Axolotl limb regeneration is accompanied by the transient induction of cellular senescence within the blastema, the structure which nucleates regeneration. The precise role of this blastemal senescent cell (bSC) population, however, remains unknown. Here, through a combination of gain- and loss-of-function assays, we elucidate the functions and molecular features of cellular senescence *in vivo*. We demonstrate that cellular senescence plays a positive role during axolotl regeneration, by creating a pro-proliferative niche that supports progenitor cell expansion and blastema outgrowth. Senescent cells impact on their microenvironment via Wnt pathway modulation. Further, we uncover a link between Wnt signalling and senescence induction, and propose that bSC-derived Wnt signals facilitate the proliferation of neighbouring cells in part by preventing their induction into senescence. This work defines the roles of cellular senescence in regeneration of complex structures.

## INTRODUCTION

The capacity for regeneration is widespread across the animal kingdom. Amongst vertebrates, urodele amphibians, such as the Mexican axolotl (*Ambystoma mexicanum*), are exceptional. Beyond the limited regenerative abilities found in mammals, they are able to regrow a broad repertoire of body parts, including their limbs, spinal cord, ocular tissues, and portions of the heart and brain^1-3^. Limb regeneration proceeds through the formation of a blastema, a collection of mesenchymal progenitor cells which accumulate at the end of the remaining stump. The blastema goes on to expand, differentiate and pattern to reconstitute a functional limb, faithfully restoring lost form and function^2,3^. Previous work by our group found that limb regeneration is accompanied by the induction of cellular senescence within a subset of cells within the blastema and surrounding tissues, referred to here as the blastemal senescent cell (bSC) population^4^.

Cellular senescence is a state of permanent growth arrest, during which cells acquire a characteristic set of phenotypic alterations including expansion of lysosomal networks, activation of tumour suppressor pathways, resistance to apoptosis, and the production of secreted factors collectively known as the senescence-associated secretory phenotype (SASP)^5-8^. The term senescence, however, does not describe a single cell state, but rather encompasses a heterogeneous set of phenotypes that vary depending on the initial triggering signal, duration of induction and tissue context^9^. Accordingly, senescence has been functionally implicated in diverse and often contradictory biological processes, mediated by a combination of cell-intrinsic growth arrest and the capacity of senescent cells to actively modulate their surrounding microenvironment through the SASP^10^. Research across the past years has established roles for senescence in a number of beneficial processes, including tissue repair^11-15^, wound healing^16,17^, embryonic development^18-21^, and cell-intrinsic tumour suppression^22^. Most beneficial forms of senescence are characterised by their transient nature, often controlled by resolution-coupled clearance, and direct or indirect tissue remodelling capacity^23,24^. Dysregulation in any of these aspects can lead to the inappropriate activity of senescent cells, driving their causal roles in pathological processes such as tissue deterioration, organismal ageing and age-related diseases^23^.

Senescent cells appear within the blastema at the early stages of regeneration, coinciding with the initiation of blastema outgrowth, and are later eliminated during the patterning phase in a macrophage-dependent mechanism^4^. The turnover of senescent cells in the blastema occurs reproducibly and repeatedly with limb amputation, and thus represents an inherent component of the regenerative program^4^. On these bases, it has been postulated that cellular senescence may play a positive function during regeneration^4^. Indeed, treatment with the BCL-2 inhibitor ABT-263, a senolytic compound, is known to delay caudal fin regeneration in zebrafish^25^. However, the precise role of cellular senescence in epimorphic regeneration is unknown and, to date, our understanding remains mostly phenomenological. Further, our knowledge on the molecular features of cellular senescence derives primarily from *in vitro* models, with limited understanding of how senescent cells behave *in vivo*.

Here, we sought to bridge these gaps, by performing in-depth characterisation of the features and functions of cellular senescence during axolotl limb regeneration, a model in which senescence induction occurs naturally *in vivo*. Through a combination of loss- and gain-of-function assays, we find that senescent cells play a critical role during limb regeneration, by facilitating progenitor-cell expansion and blastema outgrowth. Further, through bulk and single cell transcriptomic profiling, we obtain insights into the basis of senescence growth arrest *in vivo*, and elucidate the molecular basis by which bSCs create a pro-regenerative environment to support blastema outgrowth. Together, these findings identify senescence as a key component of the regenerative program, and provide insights into the biology of cellular senescence in a physiological setting.

## RESULTS

### Expansion of the senescent cell compartment accelerates regeneration

To study functional relationships between regeneration and senescence, we first employed an *in vitro* model of DNA damage-induced senescence in the axolotl limb mesenchyme-derived AL1 cell line^26^. Etoposide-treatment followed by sustained p53-stabilisation recapitulated a range of classical senescence phenotypes^4,27^, including expansion of lysosomal and mitochondrial networks, the induction of DNA damage markers, long-term proliferative arrest, and senescence-associated beta-galactosidase activity (SAβG) (Figure 1A).

**Figure 1.**
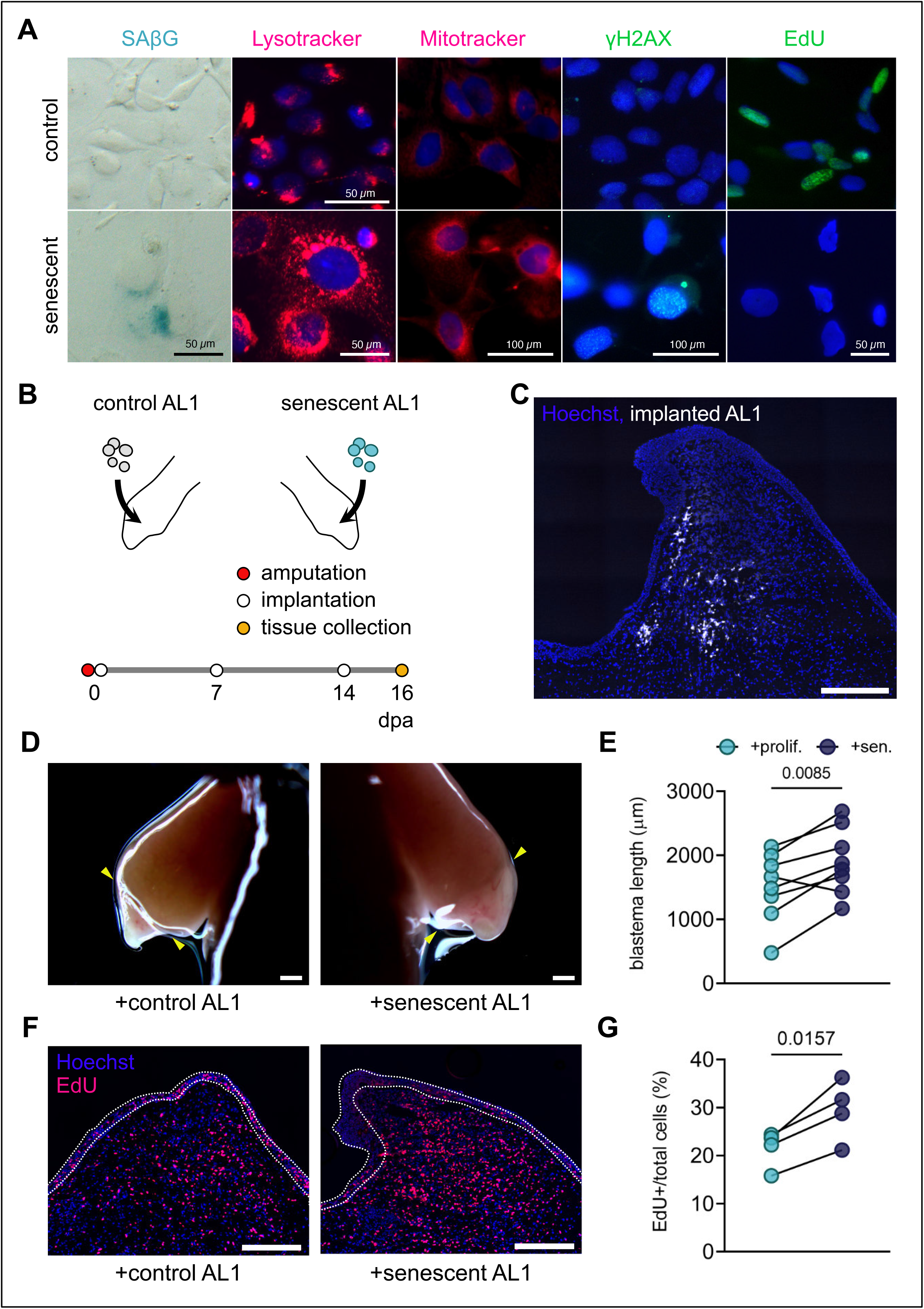
Implantation of senescent cells accelerates regeneration. **(A)** *In vitro* model of senescence in the axolotl AL1 cell line recapitulates key molecular features of cellular senescence. Proliferating control (top row) or senescent (bottom row) AL1 cells assessed for different features as indicated. Scale bars correspond to the images in each column, except for column two, as stated. **(B)** Experimental timeline for AL1 implantation studies. Proliferating and senescent cells were implanted into contralateral limbs of individual animals. **(C)** Representative longitudinal sections of 16 dpa blastema implanted with senescent AL1 cells labelled with Vybrant CM DiI. Scale bar 500 µm. Blue: Hoechst-stained nuclei; white: Vybrant CM DiI. **(D)** Representative bright-field images of contralateral limb blastemas from an individual axolotl implanted with proliferating (left) or senescent (right) AL1 cells. Scale bar = 1000 μm. **(E)** Quantification of blastema lengths at 16 dpa. P values determined by paired two-tailed t-test. Error bars depict mean ± SEM (n=8). **(F)** Senescent cell implantation stimulates cellular proliferation within the blastema. Representative sections of proliferating (left) or senescent (right) AL1-implanted blastemas stained for EdU. Scale bar = 500 μm. Blue: Hoechst-stained nuclei; magenta: EdU. White dotted lines demarcate wound epithelum. **(G)** Quantification of EdU+/total nuclei within blastema sections. P values determined by paired two-tailed t-test. Error bars depict mean ± SEM (n=4).

We began by asking whether increasing the number of senescent cells in the blastema would enhance limb regeneration (Figure 1B). Deploying a gain-of-function assay, we found that the implantation of senescent AL1 cells into regenerating tissues significantly accelerated blastema growth compared with the implantation of proliferating AL1 cells (Figures 1C-E and S1A-B), whereas the implantation of dead cells had no effect on the rate of limb regeneration (Figure S1C). Senescence-associated effects occurred specifically in the mid/late-bud phases of regeneration (Figure S1A-B), periods which demand extensive proliferation of the progenitor pool to support blastema outgrowth, prior to differentiation and patterning^28^. On this basis, we hypothesized that this enhancement of regeneration may be mediated by a senescence-associated stimulation of cellular proliferation. We found that both co-culture with senescent cells and incubation with senescent-derived conditioned media elevated EdU incorporation in AL1 cells in culture (Figure S1D-G), consistent with the production of a secreted mitogenic factor (Figures S1D-E). Furthermore, histological analysis revealed significantly increased EdU incorporation in blastemas implanted with senescent AL1 cells compared with implantation with proliferating controls, thereby demonstrating that senescence-associated effects on cellular proliferation extends to the *in vivo* context (Figures 1F-G). Given that senescent AL1 cells maintained growth arrest following implantation (Figure S1H), their *in vivo* functions are likely mediated through non-cell-autonomous mechanisms. Together, these data demonstrate that senescent cells are capable of promoting paracrine proliferation and accelerating limb regeneration in the axolotl.

### The endogenous senescent cell population supports blastema outgrowth

Next, we sought to address the role of the endogenous bSC population that arises naturally after limb amputation^4^. To do so, we first deployed a galactose-coated nanoparticle-based system, which enables selective targeting of senescent cells^29^. Their specificity predicates on the elevated β-galactosidase activity found in senescent cells, which catalyses coat degradation and thereby cargo release selectively within senescent cells (Figure 2A). We found that nanoparticles loaded with rhodamine (GalNP-rho) specifically labelled senescent AL1 cells *in vitro*, with minimal labelling of proliferating controls (Figure S2A). Further, treatment with doxorubicin-loaded nanoparticles (GalNP-dox) resulted in selective killing of senescent AL1 cells *in vitro*, with no significant effect on the viability of proliferating control cells (Figure S2B). Extending our analysis *in vivo*, we found that treatment of regenerating animals with GalNP-rho labelled a subset of cells in the blastema showing similar distribution to SaβG+ cells (Figure S2C), and importantly, we identified cells displaying both rhodamine signal and SaβG staining, confirming that galactose-coated nanoparticles label senescent cells within the blastema (Figure 2A).

**Figure 2.**
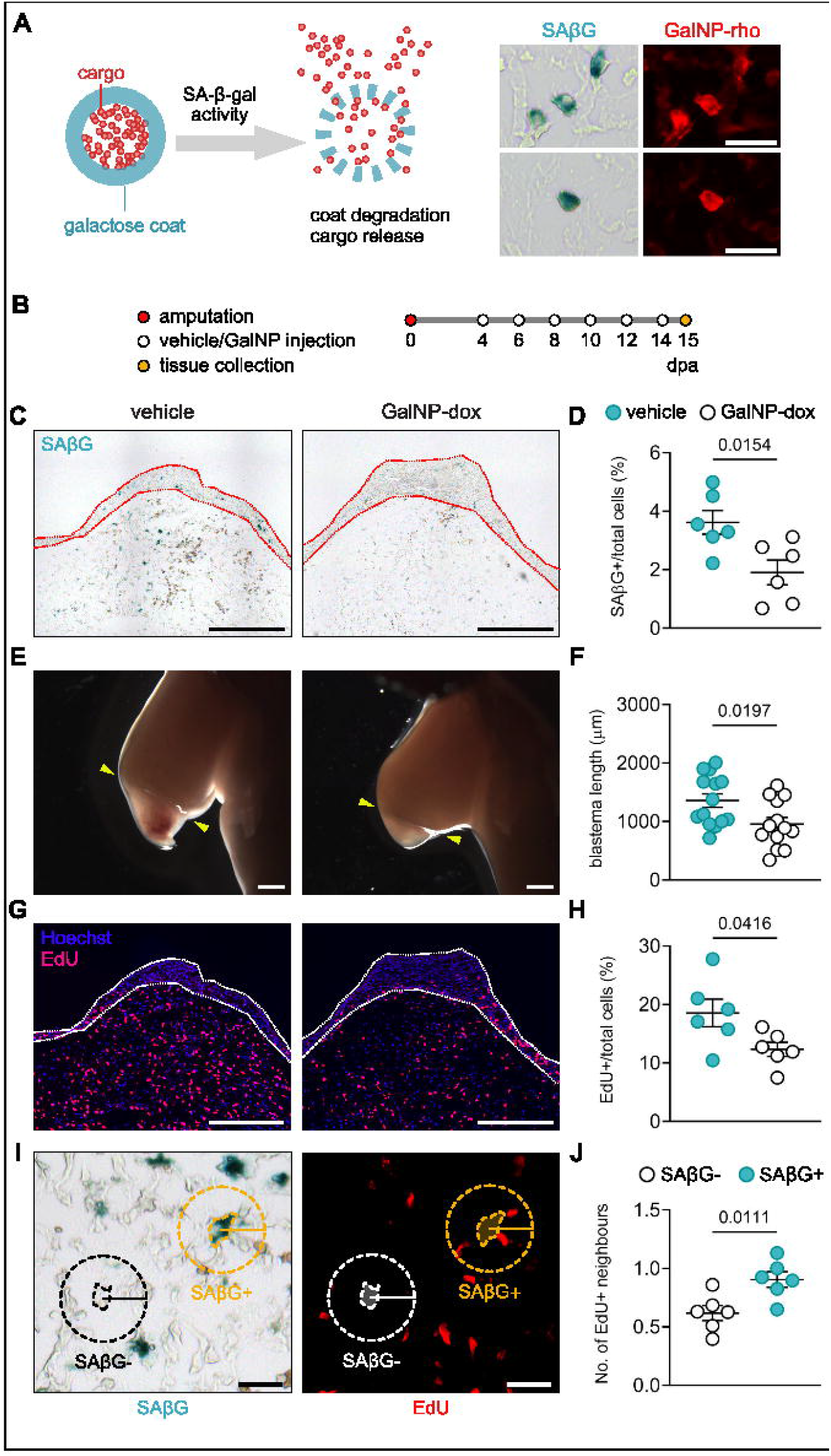
Depletion of endogenous senescent cells delays limb regeneration. **(A)** Left: schematic diagram of galactose nanoparticles. Right: dual labelling using SAβG staining and GalNP-rho in blastema sections; two examples from different sections are shown. Scale bars = 50 μm. **(B)** Experimental timeline for senescent-cell depletion studies. **(C)** GalNP-dox treatment effectively depletes the senescent population *in vivo*. Representative sections of 15 dpa limb blastemas derived from vehicle-(left) or nanoparticle-(right) treated animals with senescent cells visualised by SAβG staining (blue). Red dotted line demarcates wound epithelium. Scale bars = 500 μm. **(D)** Quantification of percentage of SAβG+/total cells within blastema sections. *P* values determined by unpaired two-tailed t-test. Error bars depict mean ± SEM (n=6). **(E)** Senescent cell depletion delays limb regeneration. Representative bright-field images of regenerating limbs from vehicle (left) or GalNP-dox (right) -treated animals at 15 dpa. Arrowheads indicate amputation site. Scale bar = 1000 μm. **(F)** Quantification of blastema length at 15 dpa. n=14 animals for vehicle and 13 animals for GalNP-dox. *P* values determined by unpaired two-tailed t-test. Error bars depict mean ± SEM. **(G)** Senescent cell depletion reduces cellular proliferation within the blastema. Representative sections (as in C) stained for EdU. White dotted line demarcates wound epithelium. Scale bar = 500 μm. Blue: Hoechst-stained nuclei; magenta: EdU. **(H)** Quantification of EdU+/total nuclei within blastema sections. *P* values determined by unpaired two-tailed t-test. Error bars depict mean ± SEM (n=6). **(I)** Neighbouring cell analysis. 15 dpa blastema sections were co-stained for SAβG (blue, left panel) and EdU (red, right panel). Average number of EdU^+^ cells surrounding senescent versus non-senescent cells within a 50 μm radius was assessed per section. **(J)** Quantification of neighbouring cell analysis. Data points represent average EdU^+^ neighbours from >3 sections analysed per animal. P values determined by paired two-tailed t-test. Error bars depict mean ± SEM (n=6).

Next, we sought to investigate the functional relevance of the bSC population towards limb regeneration (Figure 2B). GalNP-dox treatment caused significant reductions in the bSC population in regenerating tissues, as assessed by SaβG staining (Figures 2C-D), indicating that the GalNP system effectively targets senescent cells *in vivo*. Strikingly, GalNP-dox treated animals showed attenuated blastema growth, from the early- to late-bud phases of regeneration, consistent with observations from the implantation assays (Figures 2E-F and S2D-E). Assessment of EdU-incorporation revealed that GalNP-dox-treated animals displayed significant decreases in the overall level of proliferation, indicating that cellular senescence is required for maintaining a pro-proliferative microenvironment within the blastema (Figures 2G-H). In an alternative experimental paradigm, use of different senolytics (van Deursen, 2019) recapitulated delays observed with GalNP-mediated bSC depletion, consistently resulting in a ∼30% reduction in blastema length at 15 days post-amputation (dpa) compared to vehicle-treated animals (Figures S3F-I).

As senescent cells can operate in a paracrine manner^6^, we asked if the pro-proliferative effects associated with bSCs occur within their immediate microenvironment. By measuring the proliferation of nearby neighbouring cells (Figure 2I), we found that cells within a 50 μm radius surrounding senescent cells showed higher levels of EdU incorporation, as compared with those surrounding non-senescent cells, for both endogenous bSCs and implanted senescent cells (Figures 2I-J and S1C-D). Thus, these data establish that cellular senescence plays positive roles in axolotl limb regeneration and show that endogenous bSCs are able to facilitate neighbouring-cell proliferation to support blastema outgrowth.

### Transcriptomic profiling unveils the molecular basis of senescence-mediated cell-cycle arrest *in vitro* and *in vivo*

To better understand the molecular characteristics underlying senescent cells in salamanders, we next examined their transcriptomic profiles. RNA-seq analysis of proliferating and senescent AL1 cells revealed a substantial number of transcriptional changes following damage-induction of senescence (Figure S3A, Table S1). In particular, senescent AL1 cells were enriched in expression of p53-transcriptional targets, including the tumour suppressor *cdkn1a*, and displayed a corresponding reduction in the expression of E2F-target genes (Figure S3B). Additionally, senescent AL1 cells exhibited signatures of lysosomal expansion and a strong pro-inflammatory phenotype (Figure S3).

Next, we profiled endogenous bSCs and their non-senescent counterparts, purified by FACS on the basis of GalNP-rho labelling (Rho+) (Figures 3A-B, Table S2). Across regeneration stages, bSCs differentially express a number of senescence-associated markers including *glb1* (encoding SaβG), lamin B1, *oxr1* and *mmp3/10b* (Figure 3C), and display lysosomal expansion and cell cycle arrest signatures (Figures 3D and S3B-C, Table S3). In contrast to damage-induced AL1 senescent cells, endogenous bSCs are not enriched in pro-inflammatory molecules, nor do they exhibit a p53-induced damage response (Figure S3D, E). Comparison of commonly upregulated genes from *in vitro* and *in vivo* senescent cells revealed 284 transcripts shared between the two conditions (Figure 3E, Table S3). Strikingly, 16/284 overlapping transcripts correspond to pre-processed ribosomal RNA (pre-rRNA). In eukaryotes, rRNA is transcribed as a polycistronic molecule, which is cleaved and processed to yield three mature products (18S, 5.8S, and 28S) with the accompanying loss of internal and external spacer sequences (ITS1/2, ETS1/2) (Figure 3F)^30^. Notably, it has recently been reported that disruption of ribosomal biogenesis and rRNA-processing is a hallmark of senescent cells^31-33^. Gene set enrichment analysis (GSEA) revealed that genes associated with ribosomal biogenesis were significantly downregulated in both senescent AL1 and bSCs (Figures 3G and S3C), and expression of ribosomal rRNAs formed a stable signature for senescent cells across regeneration stages (Figures 3H and S3G-H). Further, reads mapping to unprocessed spacer sequences are overrepresented specifically in bSCs (Figures 3H-I and S3I). Using primers targeting 5’ ETS1 and mature 18S rRNA, quantitative PCR analysis confirmed that bSCs showed significantly higher unprocessed to processed rRNA ratios compared to their non-senescent counterparts (Figure S3J). Thus, loss of rRNA processing and ribosomal biogenesis form defining features for axolotl senescence both *in vitro* and *in vivo*. Mechanistically, disruption of these processes causes an accumulation of ribosomal proteins that no longer assemble with processed rRNA and interfere with other events^31^. In this context, Rps14 accumulation inhibits CDK4 and cell-cycle progression, inducing growth arrest in senescent cells^31^. Consistently, analysis of E2F-target genes shows a marked downregulation of cell-cycle progression genes, indicating a strong growth arrest in bSCs (Figure 3D). Furthermore, experimental overexpression of axolotl Rps14 in AL1 cells resulted in a strong reduction in cellular proliferation and an induction of SaβG activity *in vitro* (Figure S3K-L), demonstrating that phenocopying loss of ribosomal homeostasis is sufficient to induce a senescent state in salamander cells. In this connection, we note that ribosomal biogenesis processes are upregulated during early stages of limb regeneration (Figure S3M, Table S3), a significant observation given that upregulation of ribosomal proteins can elicit ribosomal stress^34^.

**Figure 3.**
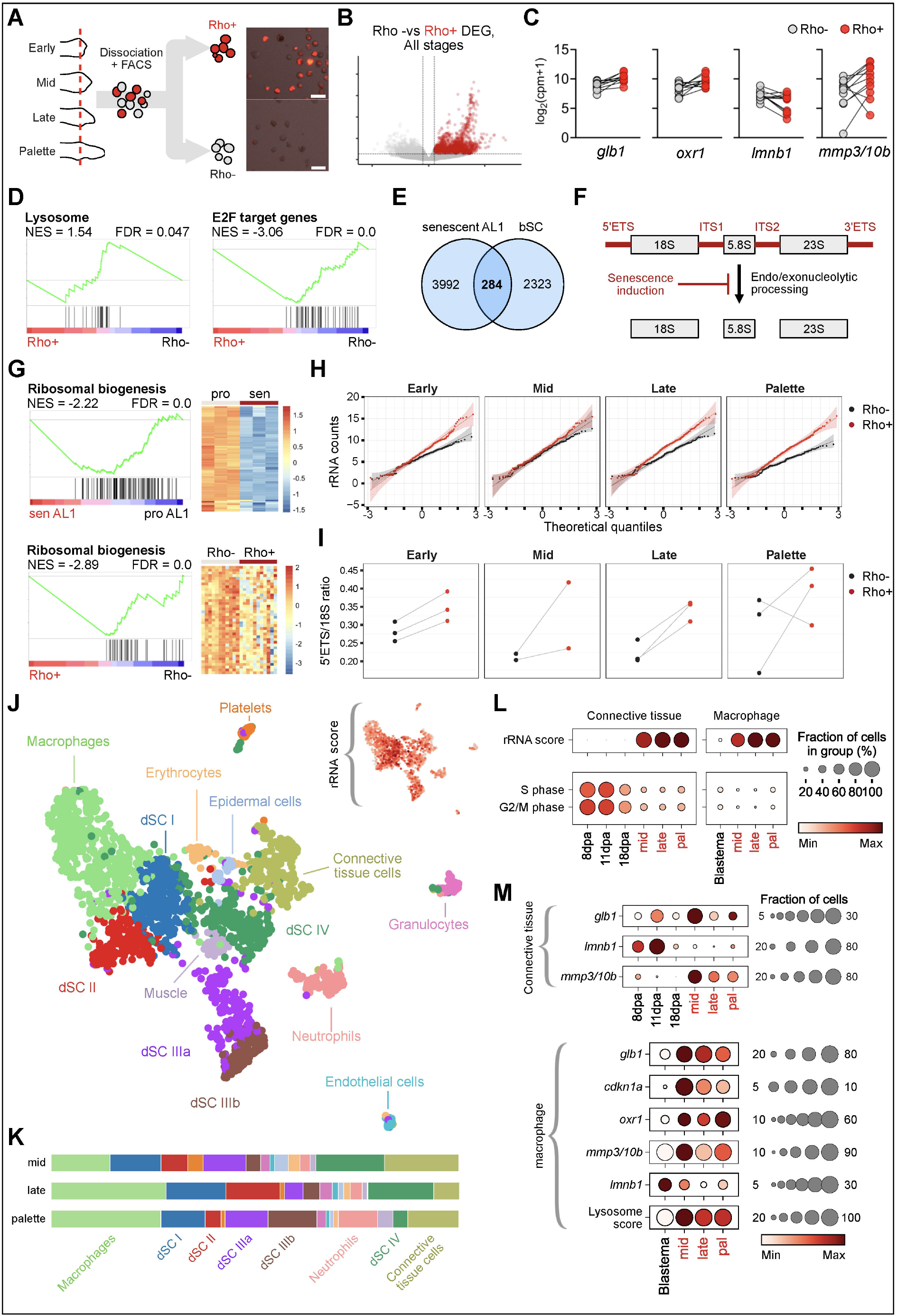
Transcriptomic profiling unveils the molecular and cellular features of bSC population. **(A)** Strategy for isolating senescent and non-senescent cells from different stages of regeneration. Insets show sorted Rho-(non-senescent) and Rho+ (senescent) blastemal cells. Red, rhodamine signal. Scale bar = 50 μm. **(B)** Volcano plot displaying differentially expressed genes between senescent (Rho+) and non-senescent cells (Rho-) isolated from different regeneration stages. **(C)** Normalized counts for selected senescence markers. **(D)** GSEA plots for lysosome and E2F target genes for blastemal senescent and non-senescent cells. **(E)** Venn diagram of commonly upregulated genes shared between *in vitro* senescent AL1 cells and endogenous bSCs. **(F)** Schematic representation of rRNA processing and its disruption in senescent cells. **(G)** GSEA plots for ribosomal processing and corresponding leading-edge analysis for *in vitro* (top) and in vivo (bottom) senescent and non-senescent cells. **(H)** Normalized counts mapping to 18S, 5’ETS, ITS1, and ITS2 ribosomal regions across indicated regeneration stages. See Supplementary Fig. 5. **(I)** Ratio of 5’ETS versus 18S transcripts in Rho- and Rho+ populations at the indicated regeneration stages. **(J)** UMAP of scRNA-seq data for individual Rho+ cells isolated from mid, late, and palette-stages of regeneration. **(K)** Corresponding cell-type proportions for (J) per regeneration stage. Marker genes for cluster identification presented in Table S4. Macrophages comprise two subclusters with increasing levels of rRNA transcripts (Table S4). **(L-M)** Differential gene expression comparisons at the single-cell level for rRNA genes and cell-cycle progression **(L)** or senescence markers **(M)**. For single-cell level comparisons, connective tissue or macrophage populations were subsetted and integrated with total connective tissue (‘8, 11, 18 dpa’, (Gerber et al., 2018)) or total macrophage (‘blastema’) single-cell datasets, respectively.

Through single cell RNA-seq analysis of purified Rho+ cells, we next sought to elucidate the population heterogeneity underlying cellular senescence *in vivo*. Gene expression-based clustering revealed that bSCs are composed of distinct subpopulations, with predominant contributions from connective tissue cells and macrophages (Figures 3J-K and S3N-O; Table S4). Importantly, transcriptional signatures of bSCs identified from analysis of bulk RNA-seq datasets remained conserved at the single-cell level independent of cell-ontogeny, as illustrated by cell-type specific comparisons between senescent cells and their non-senescent counterparts^35^ (Figures 3L-M and S3P-Q). This conserved senescence signature includes strong downregulation of G2/S/M genes, upregulation of ribosomal RNA signatures (Figure 3L) and MMP3/10b (Figure 3M). In addition, the bSC macrophage compartment displays upregulation of *oxr1, cdkn1a, glb1* and an increased lysosomal expansion signature compared to non-senescent macrophages (Figure 3M). Further, unbiased clustering revealed bSC sub-populations that lack cell-type markers, and instead express extremely high levels of ribosomal RNA (Figures 3J and S3N; Table S4), that we designated as deep-senescent clusters (dSC). Interestingly, there appears to be a transition from initial cell type to the deeply senescent state, suggested by the graded increasing expression of ribosomal transcripts from the initial cell-type into the overlapping deep-senescence clusters (Figures 3J and S3N).

### Wnt signalling is involved in senescent cell effects on their microenvironment

We next aimed to identify candidate factors responsible for bSC-mediated effects for experimental analysis, based on our functional studies, that fulfilled the criteria of being: (1) specifically enriched in senescent cells, (2) secreted, and (3) mitogenic. On this basis, we identified members of the Wnt family of extracellular ligands as potential mediators of senescence-associated effects, as they showed specific upregulation in senescent cells. In particular, *wnt7b* and *wnt8b* expression is significantly upregulated in bSCs, both at bulk and single-cell levels (Figures 4A-B). We also noted that *wnt11* and *wnt9a* are upregulated in senescent AL1 cells (Table S1). Wnts function as secreted factors that stimulate proliferation of target cells^36^ and have previously been linked to cellular senescence^37^. To experimentally address the role of Wnt signalling in senescence-derived effects, we first examined nuclear translocation of β-catenin, a key event downstream following Wnt ligand-binding that is crucial for transcriptional activation of target genes in recipient cells (Figures 4D and S4A-C)^36^. Immunohistochemical analysis revealed that senescent-cell depletion resulted in a significant reduction in β-catenin nuclear translocation in the blastema (Figure 4E). Additionally, quantitative real-time PCR (qRT-PCR) analysis revealed that expression of Wnt target genes *axin2* and *mycn* was significantly decreased in bSC-depleted tissues (Figure 4C). Thus, bSC depletion results in decreases in Wnt signalling.

**Figure 4.**
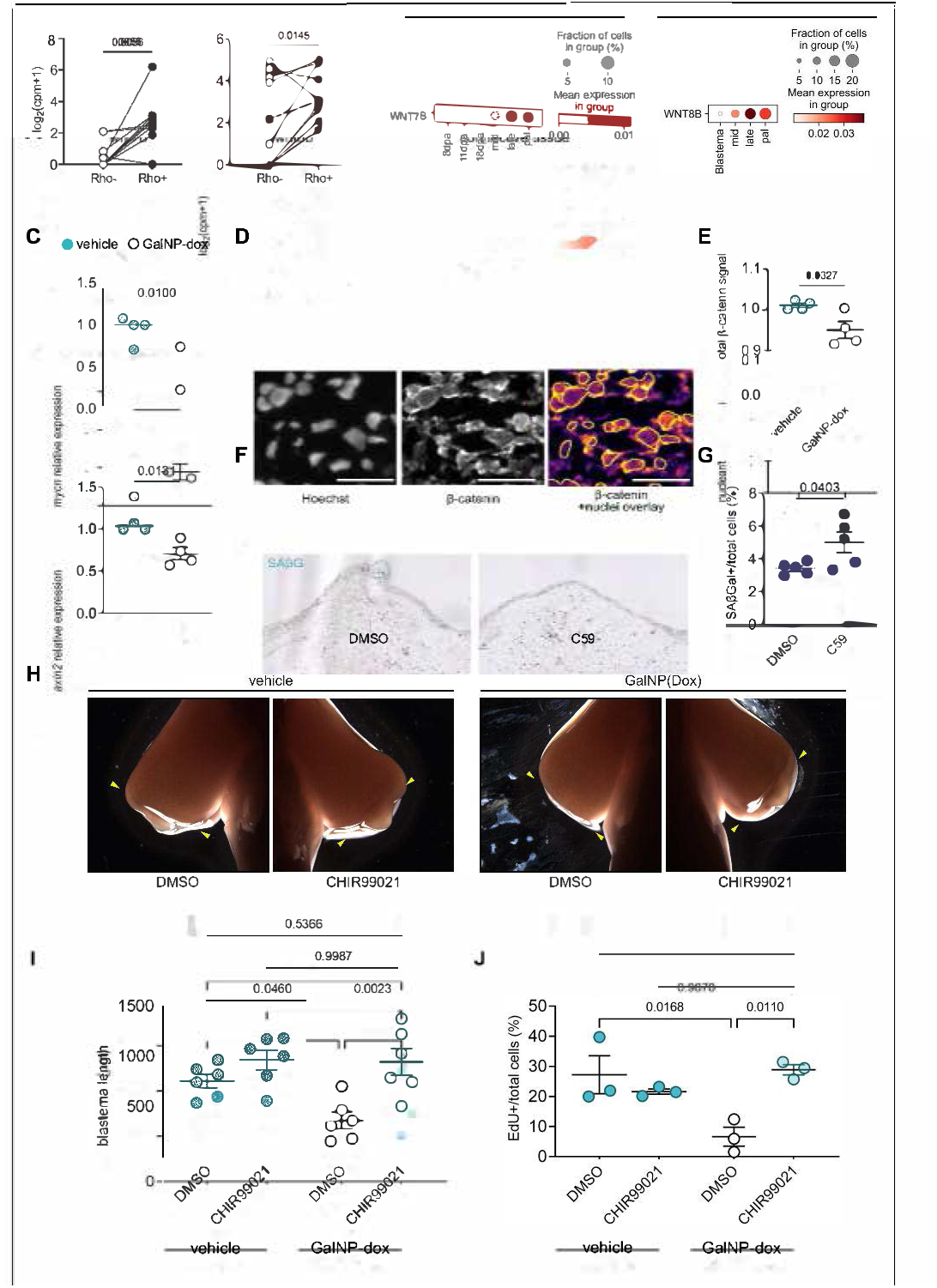
Senescent cells facilitate progenitor cell expansion through the provision of Wnt ligands. **(A)** *wnt7b* and *wnt8b* expression in bSCs and non-senescent cells measured by bulk RNA-seq. Data plotted as EdgeR normalised counts. Lines indicate pair-matched cells sorted from the same animal. **(B)** Comparison of *wnt7b* and *wnt8b* expression between total cells versus senescent (Rho+)-sorted cells at the single-cell level. For single-cell level comparisons, senescent connective tissue or senescent macrophage datasets were integrated with total connective tissue^35^ or total macrophage single-cell datasets. **(C)** mRNA levels of Wnt target genes in vehicle- or GalNP-treated blastemas measured via qRT-PCR. *axin2* and *mycn* expression values calculated using the cycle threshold method (ΔΔCT) normalised against large ribosomal protein 4 (*Rpl*4). *P* values determined by unpaired two-tailed t-test. Error bars depict mean ± SEM. **(D)** β-catenin nuclear translocation analysis. Left: Hoechst-stained nuclei. Centre: β-catenin immunostaining. Right: β-catenin immunostaining with yellow outline showing nuclear segmentation derived from Hoechst staining. **(E)** Quantification of nuclear translocation comparing vehicle- and GalNP-dox-treated blastemas, calculated by dividing mean nuclear by mean total beta-catenin signal intensity. Data points represent average values from >3 sections analysed per animal. P values determined by unpaired two-tailed t-test. Error bars depict mean ± SEM. **(F)** Representative sections of DMSO or C59-treated blastemas stained for SAβG staining (blue). Scale bars = 500 μm. **(G)** Quantification of percentage of SAβG+/total cells within blastema sections. *P* values determined by unpaired two-tailed t-test. Error bars depict mean ± SEM (n=5). **(H)** Bright-field images of regenerating limbs showing inhibition of regeneration by GalNP-dox treatment and rescue by pharmacological activation of Wnt signalling with CHIR99021. **(I)** Quantification of blastema length. P values determined by one-way ANOVA with Tukey’s multiple comparisons. Error bars depict mean ± SEM (n=6). **(J)** Quantification of EdU+/total nuclei within blastema sections showing inhibition of blastema proliferation by GalNP-dox treatment and rescue by pharmacological activation of Wnt signalling. *P* values determined by one-way ANOVA with Tukey’s multiple comparisons (n=3 blastemas per group).

We next tested whether the neighbouring-cell effect observed for bSCs (Figures 2I-J) was dependent on Wnt signalling, by treating regenerating animals with the Wnt secretion inhibitor C59 (Figure S4D). Neighbouring cell analysis on DMSO and C59-treated blastemas revealed that this neighbouring cell effect was abrogated specifically following Wnt inhibition (Figure S4E). This short-ranged effect is consistent with the post-translational attachment of a palmitoyl group to Wnt ligands, which limits their extracellular diffusion^38^. Unexpectedly, pharmacological inhibition of Wnt signalling also caused an increase in the number of senescent cells in the blastema (Figures 4F-G), suggesting that the Wnt pathway itself may limit paracrine senescence. Additionally, histochemical analysis showed that C59 treatment also resulted in increased macrophage levels (Figure S4F), which have been previously found to be recruited to senescent cells in the axolotl.

Lastly, we asked whether pharmacological activation of Wnt signalling could restore regeneration in a senescence-depleted background. Localised treatment of blastema tissue with the GSK3 inhibitor CHIR99021 effectively activated Wnt target gene expression *in vivo* (Figure S4G). In contrast to Wnt inhibition (Figure 4F-G), Wnt activation did not alter senescent cell levels in the blastema (Figure S4H). Importantly, we found that Wnt activation was sufficient to rescue limb regeneration defects caused by senescent cell depletion, as measured by EdU incorporation and blastema outgrowth (Figures 4H-J). Thus, Wnt signalling represents an important component of senescence-derived responses in regeneration.

## DISCUSSION

Here, we present a comprehensive study of cellular senescence during regeneration of complex structures. By adapting and developing methods to study cellular senescence in the axolotl model, we find that cellular senescence is required for axolotl limb regeneration and demonstrate that bSCs support the proliferation of progenitor cells in a non-cell autonomous manner, facilitating blastema expansion *in vivo* (Figure S4N). As bSCs are transiently induced following amputation, and their induction is spatially limited to damaged tissues, these functions are temporally and spatially restricted to the context of regeneration (Figure S4N).

Our transcriptomic analysis identifies key features of senescence in regeneration. First, our results suggest that growth arrest in bSC is mediated by disrupted ribosomal biogenesis, independently from activation of p53-dependent tumour suppressors (Figures 3 and S3). The disruption of ribosomal biogenesis is conserved across all stages of regeneration and forms a strong signature for cellular senescence both *in vitro* and *in vivo* (Figures 3G-I and S3). Moreover, we find that ribosomal stress directly leads to senescence induction *in vitro* (Figures S3K-L). As changes in ribosomal biogenesis are observed during early stages of axolotl limb regeneration (Figure S3M), it is possible that regeneration-associated adaptations to translational requirements result in ribosomal stress and induction of regenerative senescent cells. Although mechanistic links between ribosome biogenesis and senescence induction have been previously established, this represents a relatively under-explored aspect of senescence and its prevalence in other *in vivo* contexts merits further investigation. In addition, we uncover the heterogeneous subpopulations that comprise the total bSC population (Figure 3J) and find that bSCs consist predominantly of cells of connective tissue and macrophage-origins, with minor contributions from other cell types, consistent with recent reports of retention of cell-type identity following senescence induction^39,40^. As our characterizations relied on nanoparticle-based isolation, there is a possibility that macrophages could be non-specifically labelled due to their phagocytic activity^41^. However, our results indicate the acquisition of senescence phenotypes by a subset of macrophages during regeneration, consistent with recent reports^40,42^. This is supported by the expression of senescence signatures that distinguishes macrophages at the single-cell level (Figure 4M). Importantly, treatment with three different senolytics that act independently from phagocytic activity causes similar delays in regeneration (Figure S2F-I) as nanoparticle treatments (Figure 2), consistent with a senescence-specific effect rather than a macrophage-dependent effect. Further, senescent cells are found to stimulate paracrine cellular proliferation *in vitro* in the absence of macrophages (Figure S1D-G). In line with this, macrophages did not elicit localised increases in cellular proliferation in their neighbours (Figure S4I-J), as is observed for senescent cells. Importantly, the neighbouring-cell effect associated with senescent cells persisted after macrophage depletion (Figure S4K-M), suggesting that at least a subset of senescent cells (of non-macrophage origin) can themselves directly exert pro-proliferative effects.

Mechanistically, our results suggest that the pro-regenerative effects of bSCs are mediated by Wnt signalling (Figure 4). Previous links have been observed between Wnt and senescence in the context of responses to therapy in cancer^37,43^. Cells that manage to escape therapy-induced senescent growth arrest acquire stem-cell signatures, increased growth capacity and tumorigenic potential in a Wnt-dependent manner^37^. Conceptual parallels can be drawn between the regeneration and cancer contexts; both are characterized by broad, tissue level damage that results in two distinct outcomes for damage-exposed cells – either senescent growth arrest or extensive proliferation. Both scenarios can be interpreted within the general framework that senescence occurs in response to extensive tissue damage, simultaneously preventing the expansion of damaged cells whilst orchestrating repair of the surrounding tissue compartment. In addition, parallels can be drawn between bSCs and ‘undead’ cells in the fly wing disc regeneration model, in which prevention of death of apoptotic cells results in their secretion of Wnt ligands which foster compensatory proliferation^44^. Further, whether senescence-derived Wnt can impact on additional mechanisms relevant to blastema formation, such as dedifferentiation or crosstalk with other signalling pathways, is currently unknown. Interestingly, activation of canonical Wnt signalling during limb regeneration has recently been found to elicit the expression of Fgf ligands, which are essential for sustaining blastema outgrowth^45-47^. The relative contributions of senescent-specific Wnt subtypes towards this process awaits further exploration. Lastly, the possibility that Wnt-independent senescent cell functions exist remains open, given the discrepancy between senescent cell implantation and broad Wnt activation in normal conditions.

Intriguingly, broad pharmacological inhibition of Wnt signalling caused a higher burden of senescence within the blastema, suggesting the existence of feedback mechanisms (Figures 4F-G). A recent report from the Berge group showed that experimental activation of Wnt signalling facilitates mesenchymal stem cell expansion *in vitro*, not only through direct stimulation of proliferation *per se*, but also by suppressing the spread of paracrine senescence from an initial sub-population of senescent mesenchymal stem cells^48^. Of relevance, prevention of senescence transmission increases hepatocyte proliferation and improves tissue renewal following severe liver damage in murine models^49,50^. Thus, it is possible that bSCs-derived Wnts may primarily protect surrounding cells from entering senescence in the face of amputation-derived signals, and pro-proliferative effects may be secondary. In line with this view, endogenous bSCs do not exhibit a pro-inflammatory signature (Figure S3) and have decreased expression of factors implicated in paracrine senescence in mammalian and *in vitro* models^5^. Of note, activation of Notch signalling in senescent cells induces expression of the Notch ligand Jag1 upon oncogenic Ras expression, inducing senescence in neighbouring cells through a ‘bystander mechanism’^51,52^. Consistent with a senescent-suppressing activity, bSCs display significant downregulation of Notch activation (Figure S3E). Whether this results from a Wnt-Notch molecular crosstalk remains an outstanding question. Together, our findings suggest that in the axolotl, Wnt production and the lack of a pro-inflammatory phenotype collectively prevents the propagation of senescence across regenerating tissues (Figure S4N), and may represent an important difference in the senescence response to severe injury between mammals and salamanders.

This work uncovers the functions of cellular senescence during regeneration of complex structures, offering mechanistic insights into its nature and highlights the positive impact of cellular senescence in a physiological regenerative context. As such, it has significant fundamental and therapeutic implications.

## Limitations of the study

Our work provides significant evidence that senescent cells impact on progenitor cell proliferation in a short-range manner via WNT pathway modulation, however the abrogation of WNT ligand production specifically in senescent cells is not technically feasible at present. Advances in selective knock-out/in/down in senescent cells, e.g. through genetic or nanoparticle-based strategies, will hopefully provide in-depth mechanistic insights in the future. Further, while we demonstrate that GalNP-dox nanoparticles selectively eliminate senescent cells, the specific killing mechanism remains to be addressed.

## Supporting information

Supplemental material combined

## ACKNOWLEDGMENTS

We thank TUD-CMCB Flow Cytometry and Light Microscopy Facilities for cell sorting and imaging support, DRESDEN-concept genome center for RNA sequencing, MPI-CBG Computing Facility for image processing advice, Phillip Gates for technical support, Daniel Muñoz Espin for nanoparticle advice, Gabriel Waksman for institutional support, Beate Gruhl, Anja Wagner and Dominic Kruger for animal care, and all members of the Yun lab for advice and comments on the manuscript.

## AUTHOR CONTRIBUTIONS

QY, HEW, PK, ML, CRO, CSA, DK, AF, ABB, MAA and MHY designed and performed experiments; QY, HEW, GP, SPS, DLP, AP, PK, IG, CSA, DK, AF, ABB, MAA, GN, AMG, KET, AD, VB, RMM and MHY analysed and interpreted data. QY and MHY wrote the manuscript with input from all authors. MHY supervised the study.

## STAR METHODS

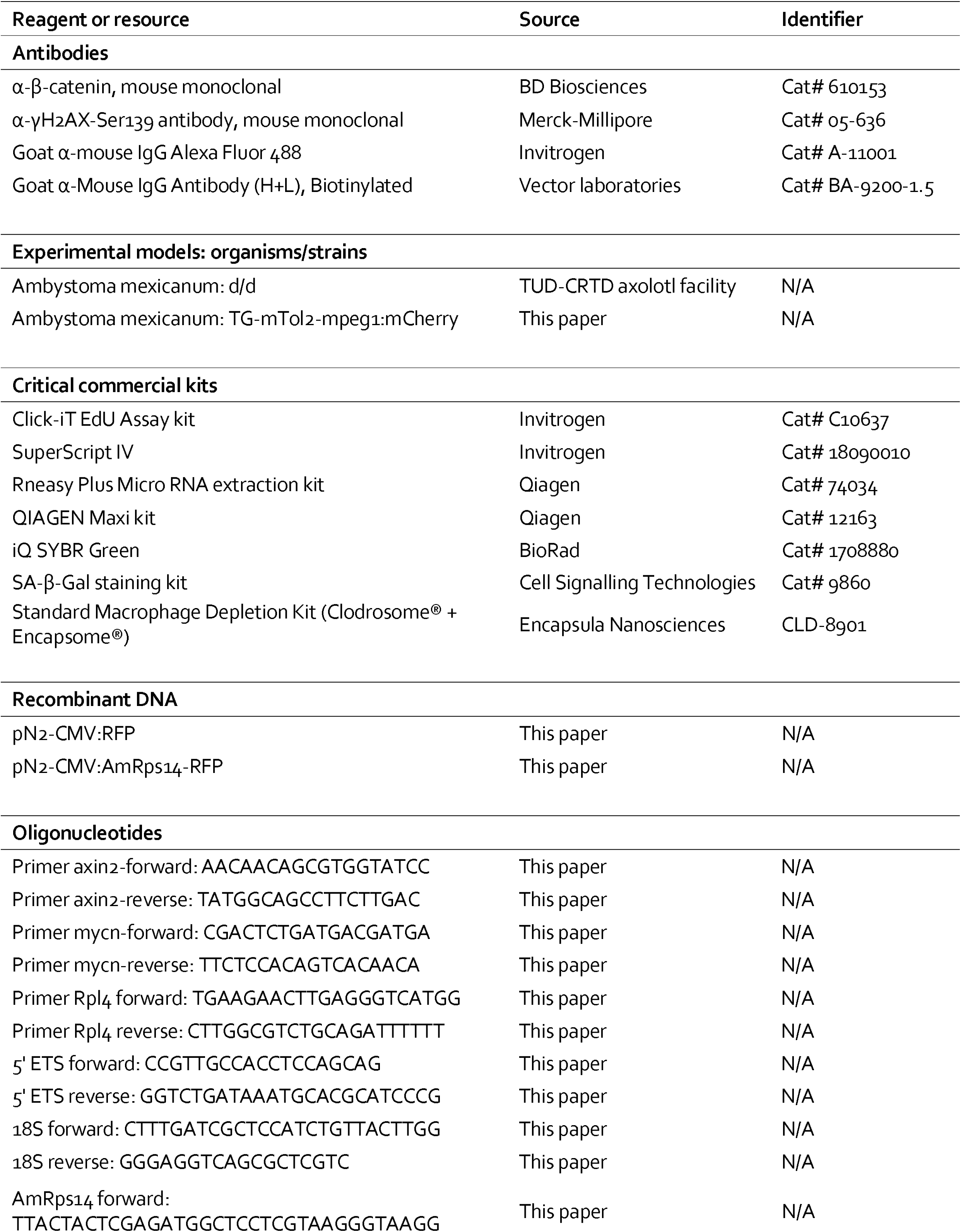
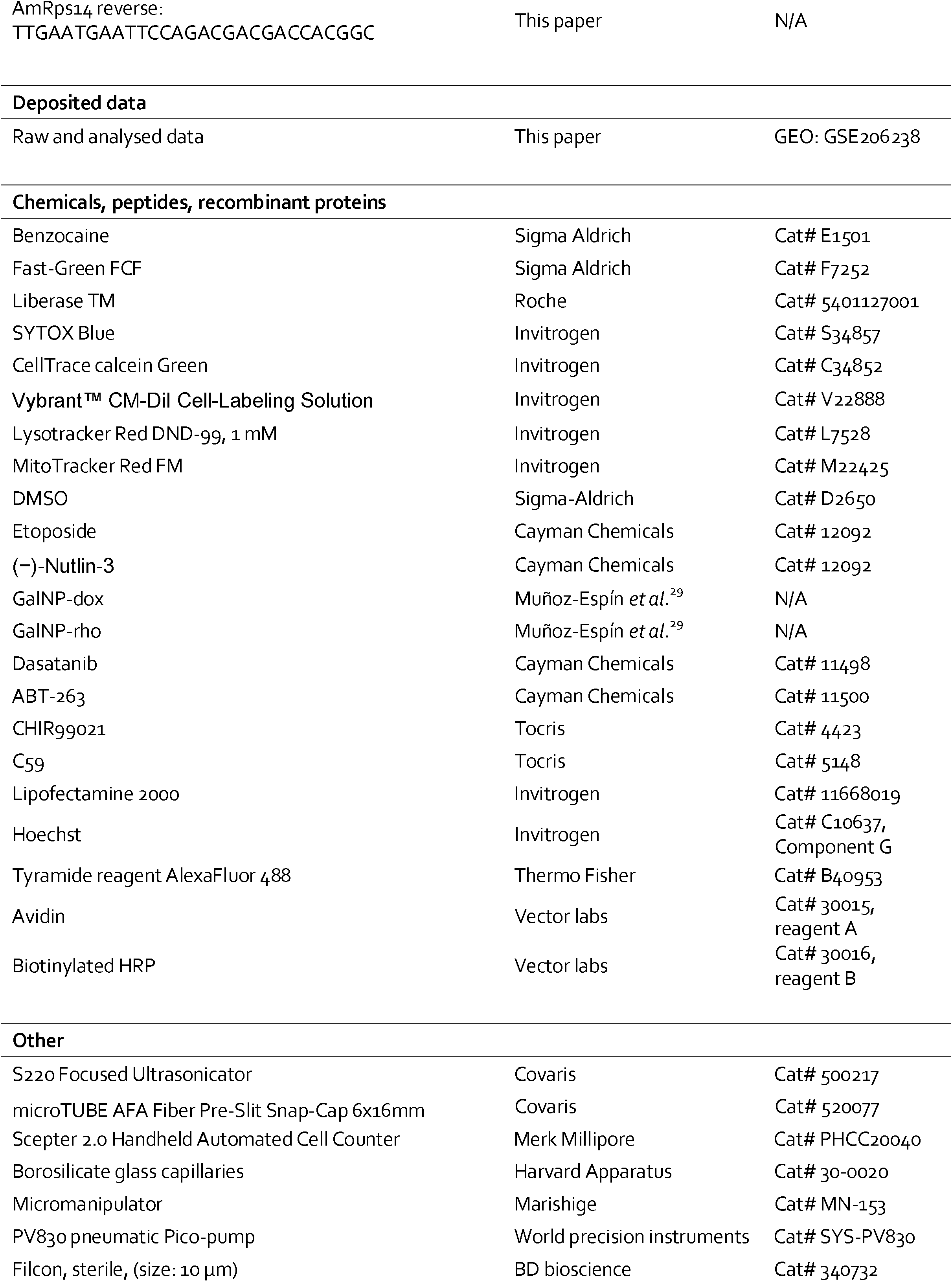

## RESOURCE AVAILABILITY

### Lead contact

Further information and requests for resources and reagents should be directed to and will be fulfilled by the Lead Contact, Maximina H. Yun.

### Materials availability

Academic labs will have access for non-profit research with Material Transfer Agreement.

### Data and code availability

- All bulk and single-cell RNA-seq data have been deposited at GEO and are publicly available as of the date of publication. Accession numbers are listed in the key resources table. Microscopy data reported in this paper will be shared by the lead contact upon request.
- All original code has been deposited at https://github.com/yun-crtd/Yu-et-al-2022 and is publicly available as of the date of publication.
- Any additional information required to reanalyze the data reported in this paper is available from the Lead Contact upon request.

## EXPERIMENTAL MODEL AND SUBJECT DETAILS

### Axolotl strains and maintenance

Axolotls (*A. mexicanum*) were obtained from Neil Hardy Aquatica (Croydon, UK) and from the axolotl facility at TUD-CRTD Center for Regenerative Therapies Dresden (Dresden, Germany). Axolotls of the leucistic (d/d) strain were used in all experiments. Juvenile (6-20 cm in size) and mature (over 21 cm) axolotls were used in this study. All animal procedures used in this study were performed in compliance with the Animals -Scientific Procedures-Act 1986 (United Kingdom Home Office), and the laws and regulations of the State of Saxony, Germany.

#### Husbandry conditions

Animals were maintained in individual aquaria at ∼18–20 °C, as previously described^53^. Housing space is adjusted to animal size according to the regulations of the state of Saxony. Juveniles were daily fed small axolotl pellets (3 mm, Axolotlpellets AXOBALANCE, Aquaterratec), while adults were fed thrice a week with protein-rich axolotl pellets (4.5 mm, Axolotlpellets AXOBALANCE, Aquaterratec).

## METHOD DETAILS

### Surgical procedures

For all animal procedures, axolotls were anaesthetized in 0.03% benzocaine (Sigma), and procedures performed under an Olympus SZX10 stereoscope.

### Amputations

Axolotls were amputated through the upper humerus, and allowed to regenerate at 20°C.

### Microinjections

Micro-injections described in this study were performed using borosilicate glass capillaries (Harvard Apparatus, cat. no. 30-0020) pulled with a Flaming/Brown micropipette puller P-97 (Sutter Instrument Company). Capillaries were pulled to generate long-tapered and thin-bored tips to minimize tissue disruption during injections. Injections were performed with the aid of a micromanipulator (Science Products, Marishige MN-153) and pressure controlled by a PV830 pneumatic Pico pump (WPI). Solutions were supplemented with 0.2 mg ml^-1^ Fast Green-FCF (Sigma Aldrich) to visualise injections.

### Cell culture

AL1 cells, initially derived by S. Roy (University of Montreal, Montreal, Canada), were grown on 0.75% gelatin-coated plastic dishes in MEM (Gibco, UK) complemented with 10% heat-inactivated foetal calf serum (FCS, Gibco), 25% H_2_O, 2⍰nM L-Glutamine (Gibco), 10⍰μg/ml insulin (Sigma, St Louis, MO) and 100⍰U/ml penicillin/streptomycin (Gibco) in a humidified atmosphere of 2.5% CO_2_ at 25⍰°C. Cell culture was performed as described (Denis et al., 2015).

Conditioned media was generated by exposing fresh complete media to proliferating or senescent AL1 cultures at comparable confluence for 48 hours, prior to filtration (0.22⍰μm). AL1 cells were then treated with freshly-generated conditioned media for 24 hours, with a 5 uM EdU pulse for the final 4 hours prior to fixation to quantify proliferation.

For co-culture experiments, plates of senescent or proliferating AL1 cells were stained with CellTracker Green according to manufacturer’s instructions, and washed three times with A-PBS. Cells were then harvested with Trypsin, and seeded at equal density in co-culture (with one stained and one control un-stained population respectively). Cells were incubated for 24 hours, with a 5 uM EdU pulse for the final 4 hours prior to fixation to quantify proliferation in stained or unstained populations.

### AL1 cell implantation experiments

#### Senescence induction

*In vitro* senescence induction was performed according to^27^, whereby cells were seeded at 50% confluence and subsequently exposed to 10 μM etoposide and 1 μM nutlin-3a for 24h. Cells were then incubated with 1 μM nutlin-3a in fresh media for the following 11 days. Control proliferating cells were seeded in parallel, treated with identical volumes of DMSO only, and passaged during the 12-day senescence induction to avoid confluence. Senescence induction was confirmed by positive SAβG staining, reduction in EdU incorporation, expansion of mitochondrial and lysosomal networks, and persistent DNA damage foci (MitoTracker, Lysomitotracker, γH2AX;^27^).

#### AL1 cell harvesting & implantation

On the day of implantation, media was exchanged for fresh growth medium supplemented with 5 μM Vybrant CM DiI dye for 30 min. Subsequently, cells were rinsed twice with 0.8X PBS, trypsinised, and density determined using a Scepter 2.0 Handheld Automated Cell Counter (Merk Millipore). Cells were then pelleted and resuspended at a density of 5000 cells/μl in 0.8X PBS/0.2 mg ml^-1^ Fast Green-FCF. 10 μl of cell suspension were locally injected into the upper limb mesenchyme. The upper limb was subsequently amputated, distally from the site of cell implantation. Implantations were repeated at 7 and 14 dpa by localised microinjection into the blastema mesenchyme. At 16 dpa, blastemas were harvested for processing and analysis.

### *In vivo* drug treatments

#### Preparation of GalNP suspensions

Nanoparticles were weighed on a microscale, and resuspended in 0.8X MEM/0.2 mg ml^-1^ Fast Green-FCF at a final concentration of 4 mg ml^-1^. Suspensions were aliquoted into S2 microfibre tubes (Covaris). Suspensions were sonicated to disperse nanoparticle electrostatic aggregates and facilitate intravenous injection through thin-bore glass capillaries, using an S2 water-bath sonicator (Covaris), under the following parameters: duty cycle, 18%; intensity, 7.5; cycles/burst, 1000; duration, 10 sec; mode, frequent sweep. Immediately after sonication, 120 μl of nanoparticle suspension were injected intravenously into gill arteries (corresponding to ∼2 μl per gram of animal weight).

#### Nanoparticle administration

For senescence depletion experiments, vehicle or GalNP-dox was administered every 48 h from 4 dpa onwards, and when indicated, blastemas harvested for analysis at 15 dpa. For GalNP-mediated labelling of senescent cells, axolotls were injected with GalNP-rho and blastemas collected 48 h after injection, dissociated, and subjected to live/dead staining and FACS as described below.

#### C59 treatments

A 5 μM solution of C59 was prepared by diluting a 5 mM stock in DMSO 1:1000 in 0.8X PBS/0.2 mg ml^-1^ FG-FCF. 30 μl was administered by intravenous injection into gill arteries at 10, 12, and 14 dpa. At 15 dpa, animals were subjected to a 3.5 h pulse of EdU as described below prior to tissue collection. Senolytic treatments. For senolytic treatments, DMSO, dasatinib, or fisetin was administered every 48 h from 4 dpa onwards. Dasatanib was delivered at 0.2 μg per gram of animal weight, and fisetin at 1 μg per gram of animal weight through IV delilvery. ABT263 was administered every 48 h, from 7 dpa onwards, at a concentration of 10 μg/g of animal weight through IP delivery.

#### CHIR99021 treatments

A 1.5 μM solution of CHIR99021 was prepared from a 1.5 mM stock in DMSO by diluting in 0.8X PBS/0.2 mg ml^-1^ FG-FCF. 2 μl was locally injected into blastema mesenchyme using glass capillaries as described above, every other day between 4 until 14 dpa, under the same treatment schedule as nanoparticle injections. At 15 dpa, animals were subjected to a 3.5 h pulse of EdU as described below prior to tissue collection.

#### Clodrosome treatments

Clodrosome treatments were performed as previously described^4^. Clodrosome and control PBS liposomes solutions were obtained from Encapsula Nanosciences Ltd. Animals were injected intravenously (20 µl solution for 17cm snout-to-tail axolotls) through gill arteries at 10, 12, and 14 dpa. At 15 dpa, animals were subjected to a 3.5 h pulse of EdU as described below before blastema collection.

#### EdU treatments

For in vivo EdU labelling, animals were injected intraperitoneally using a 30-gauge needle and syringe, using 10 μl of 0.8X PBS/0.2 mg ml^-1^ Fast Green-FCF/1 mg ml^-1^ EdU per gram of animal weight. Blastemas were then harvested at 3.5 h post-injection for histology as described above.

### Sample collection

#### RNA extraction, cDNA synthesis and qRT-PCR

For RNA extractions, samples were collected into 350 μl β - mercaptoethanol-supplemented RLT+ buffer and lysed through pipetting and manual dissociation when necessary. Total RNA was then extracted using RNeasy micro plus RNA extraction kit (Qiagen) according to manufacturer’s instructions. 800 ng of total RNA was used as template for cDNA synthesis using SuperScript IV First Strand Synthesis System (Invitrogen) and 2.5 μM random hexamers primers.

Quantitative PCR analysis of target gene expression was performed using 1x SYBR Green supermix (BioRad) and 500 nM forward and reverse primers using standard curves per run. The concentration of *axin2* and *mycn* were normalized with that of Rpl4 (large ribosomal protein 4). Primers used for qRT-PCR are shown in Key Resources Table - Oligonucleotides.

#### Cryosectioning

Samples were collected into PBS/4% PFA and fixed overnight at 4 °C. Afterwards, samples were washed for 2x 10 min in 1X PBS, embedded in optimum cutting temperature (OCT) compound in plastic moulds and frozen at -80 °C. Samples were sectioned into 8 μm thick sections and collected onto Superfrost-plus microscopy slides (Menzel-Gläser). Sections were airdried for 1 h before storing at -20 °C.

### Histology and stainings

#### SAβG and EdU stainings

SAβG and EdU stainings were performed as previously described^4^. Briefly, slides were airdried at room temperature for 45 min and rehydrated in PBS. Slides were then incubated in SAβG staining solution adjusted to pH 6.5 (Cell Signalling Technologies, prepared according to manufacturer’s instructions) for 16 h at 37 °C in a humified chamber. Subsequently, slides were washed 3x in PBS for 10 min, permeabilised in PBS/0.3% TX100 for 15 min, and blocked in PBS/0.3% Triton X100/10% goat serum for 1 h. EdU detection was then performed by incubating with EdU staining solution according to manufacturer’s instructions (Click-iT EdU Assay, ThermoFisher). Slides were counterstained with Hoechst, mounted and sealed for imaging.

#### NAE stainings and EdU stainings

Monocytes/macrophages were identified using the α-naphtyl acetate esterase (NAE) kit (Sigma–Aldrich), as previously described^4^. Note that the chemical reaction for 10-12µm sections was conducted for only 10 minutes. Following 2x washes in dH_2_O, sections were fixed in 4% PFA. NAE staining was followed by 5-ethynyl-2⍰-deoxyuridine (EdU) detection using Click-iT Edu Alexa Fluor 488 or 594 Imaging kits (Life Technologies) according to the manufacturer’s instructions.

#### β-catenin antibody stainings

Slides were airdried at room temperature for 45 min and rehydrated in PBS. Slides were then incubated in 10 mM sodium citrate/0.05% Tween, pH 6.0 for 15 min at 100 °C for antigen retrieval, and washed 3x in dH_2_O for 3 min. Slides were subsequently incubated in PBS/3% H_2_O_2_ to quench endogenous peroxidase activity. Afterwards, sections were permeabilised in PBS/0.3% Triton X100, blocked in PBS/0.3% Triton X100/10% goat serum for 1 h, and incubated with primary antibody diluted in blocking buffer overnight at 4 °C in a humidified chamber. Subsequently, slides were washed 3x in PBS/0.3% Triton X100, incubated with biotinylated secondary antibody in blocking buffer (1:200 dilution) for 1 h, and then incubated in PBS/avidin/biotinylated-HRP for 1 h. Tyramide detection was performed by incubating in AlexaFluor 488 tyramide reagent diluted 1:100 in tyramide amplification buffer (0.1% Tween-20/0.003 % H2O2/100 mM borate, pH 8.5) for 10 min at room temperature. Washes with PBS/0.3% TX100 were performed between all incubations (3x 5-min washes). Slides were counterstained with Hoechst, mounted and sealed for imaging.

### Rps14 transfection studies

#### Rps14 plasmid cloning

AmRps14 was amplified from axolotl blastema cDNA using the following primers: AmRps14_forward: 5’-TTACTACTCGAGATGGCTCCTCGTAAGGGTAAGG-3’; AmRps14_reverse: TTGAATGAATTCCAGACGACGACCACGGC. The amplicon was subcloned into pN2-CMV:RFP plasmid using XhoI/EcoRI restriction enzyme digestion-ligation (NEB) to generate pN2-CMV:AmRps14-RFP plasmids and sequence verified. Plasmids used for lipofection studies were prepared with a Qiagen Maxi kit (Qiagen).

#### Rps14 lipofection

AL1 cells were seeded into 35 mm Nunclon TM dishes at 70-80% confluence. At 24 h post-seeding, media was exchanged for 1.5 ml antibiotic free growth medium (MEM (Gibco, UK) complemented with 10% heat-inactivated foetal calf serum (FCS, Gibco), 25% H_2_O, 2⍰nM L-Glutamine (Gibco), and 10⍰μg/ml insulin (Sigma, St Louis, MO)). Transfection mixtures were prepared by mixing 8 μl of Lipofectamine 2000 (Invitrogen) with 3 μg of plasmid DNA in 400 μl Opti-MEM (Gibco), and added to each dish. Following overnight incubation, lipofection media was aspirated and exchanged for normal growth medium, and grown for 6 days in a humidified atmosphere of 2.5% CO_2_ at 25⍰°C.

#### Analysis of Rps14 overexpression on cell-cycle

For EdU experiments, cells were pulsed with 5 μM EdU for 4 h at 6 days post-transfection, and then rinsed with 0.8X PBS and fixed in PBS/4% PFA for 15 min. Subsequently, EdU detection was then performed by incubating with EdU staining solution according to manufacturer’s instructions (Click-iT EdU Assay, ThermoFisher). Dishes were counterstained with Hoechst, mounted and sealed for imaging.

#### Analysis of Rps14 overexpression on SAβG

For SAβG studies, cells were collected at 17 days post lipofection, washed in 1X PBS, fixed in 0.5% glutaraldehyde and stained for SAβG activity as described^4^.

### Imaging and image analysis

Images were acquired using a Zeiss AxioZoom V16 microscope and Zen 2.3 software (Zeiss). The same imaging conditions (exposure, magnification) were used for all samples within a given experiment.

#### SAβG, NAE and EdU quantifications

Quantifications were performed using FIJI^54^. Cell nuclei were segmented using the Otsu thresholding method, and quantified in Hoechst and EdU channels to obtain total and EdU-positive nuclei, respectively. SAβG and NAE positive cells were counted manually on the basis of the presence of SAβG or NAE signal, respectively. Data are presented the percentage of positive cells as normalised against the total number of nuclei.

#### Neighbouring cell analysis

Neighbouring cell proliferation was measured on 3.5 h EdU-pulsed blastema samples from 1 year old d/d axolotls. Blastema sections were co-stained for EdU and SAβG or NAE, as described above, and neighbouring-cell EdU incorporation was assessed using a custom Python script (https://github.com/yun-crtd/Yu-et-al-2022). Briefly, for senescent-cell neighbouring cell analysis, cells were selected as senescent or non-senescent on the basis of SAβG staining in bright-field images. EdU-positive nuclei were segmented using CellPose, and the number of EdU-positive nuclei within a 50 μm radius surrounding selected cells were quantified using the custom Python script neighborhood_sab_edu_vx.x.py (https://github.com/yun-crtd/Yu-et-al-2022). For macrophage-based neighbouring cell analysis, cells were selected as macrophage or non-macrophage on the basis of NAE signal in bright-field images, and subsequently, the number of EdU-positive nuclei within a 50 μm radius was quantified as described above.

#### Nuclear translocation quantifications

β-catenin nuclear translocation analysis was performed using FIJI^54^ on vehicle or GalNP-dox-treated blastema sections stained for β-catenin and Hoechst, as described above. Cell nuclei were segmented using the Otsu thresholding method to obtain nuclear boundaries; total β-catenin signal segmented using the Huang threshold method to obtain total β-catenin boundaries, in order to normalise for differences in tissue density across sections. Masks were overlayed on top of β-catenin channel images, and average intensity was measured within nuclear boundaries or total β-catenin signal boundaries, as indicated in Fig. S7, to obtain average nuclear and total β-catenin intensities. β-catenin nuclear translocation ratios were then calculated by dividing nuclear β-catenin intensities by total β-catenin intensities.

#### AmRps14 transfection EdU quantifications

Quantifications were performed using FIJI. Cell nuclei were segmented using the Otsu thresholding method, and quantified in Hoechst and EdU channels to obtain total and EdU-positive nuclei, respectively. Transfected cells were identified manually on the basis of RFP expression, and the number of EdU-positive and -negative nuclei then quantified.

#### AmRps14 transfection SAβG quantifications

To determine the number of total cells, nuclei were quantified using FIJI based on Hoechst staining, as described above. Senescent cells were quantified manually, based on the presence of X-gal staining (blue colour) and expressed as percentage of senescent cells/total cells.

### Transcriptomic studies

#### Tissue dissociation and FACS isolation of senescent and non-senescent blastema cells

Blastemas samples were collected from mid-, late- or palette-stage animals at 48 h following GalNP-rho injection. Samples were dissociated in 400 μl of 1X Liberase (Roche) diluted in 0.8X PBS, using mechanical disruption with forceps together with smooth pipetting at room temperature for 30 min. For live/dead staining, suspensions were incubated in 0.8X PBS supplemented with 200 nM CellTrace calcein green AM (Invitrogen) and 12.5 μM SYTOX blue (Invitrogen) at room temperature for 15 min. Subsequently, cell suspensions were filtered through a 70-μm-diameter strainer to generate single-cell suspensions and immediately subjected to FACS. Alive cells were sorted using a FACSAria III equipped with a 100 μm nozzle (BD Biosciences) according to the presence or absence of rhodamine signal using blastemas from vehicle-injected axolotls as negative controls.

For scRNA-seq experiments of bSCs, single rhodamine-positive cells were sorted into 2 μl of lysis buffer (nuclease-free water, 0.2% Triton X100, 4U/μl RNase Inhibitor) in individual wells of 396-well plates, and stored at -80 °C until further processing as described below. In the case of bulk RNA-seq analysis of bSCs, for each given animal, equivalent numbers of senescent (Rho+) and non-senescent (Rho-) cells were FACS isolated and sorted into separate PCR tubes containing 10 μl lysis buffer, and stored at -80 °C until further processing.

#### Senescent and proliferating AL1 RNA-seq sample preparation

For RNA-seq analysis of AL1 cells, senescent or proliferating control cells in 10 cm dishes were harvested at 14 days post induction. Briefly, cells were rinsed twice with 0.8X PBS, trypsinised, and pelleted. Subsequently, pellets were lysed in 350 μl β-mercaptoethanol-supplemented RLT+ buffer. Total RNA was then extracted using RNeasy micro plus RNA extraction kit (Qiagen) according to manufacturer’s instructions. 1000 ng of total RNA was used as input for library preparation as described below.

#### Library preparation

All samples were processed following the Smartseq-2 protocol (complete details are found at https://star-protocols.cell.com/protocols/1595). Briefly, samples were thawed on ice and 0.5 μl of dT-buffer was added to each sample, containing poly-dT oligos (final concentration of 0.5 μM) for capturing mRNAs. Subsequently, we performed reverse transcription and cDNA amplification. Libraries were prepared with the Vazyme TruePrep DNA Library Prep Kit V2 (bSC bulk RNA-seq samples; kit discontinued) or the Illumina® DNA Prep Tagmentation kit (bSC scRNA-seq and AL1 samples). Libraries from bSC scRNA-seq samples were sequenced on an Illumina NovaSeq 6000 S1 flow cell to an average of 0.85 mio PE50bp fragments/cell, libraries from bSC bulk RNA-seq samples on an Illumina NextSeq 500 to an average of 32mio SE75bp fragments/sample and libraries from AL1 samples on an Illumina NextSeq 6000 to an average of 48mio PE100bp fragments/sample.

#### Mapping and counts

Sequence and gene annotation of the axolotl nuclear genome assembly AmexG_v6.0-DD were downloaded from https://www.axolotl-omics.org. The gene models for the HoxA and HoxD genes were replaced with manually curated gene models (personal communication with Sergej Nowoshilow, axolotl-omics.org). Sequence and gene annotation of the mitochondrial genome assembly (NCBI GenBank, AY659991.1) and RNA spike-in control sequences (ERCC, Ambion; for quality control) were included as well. RNA-seq reads were mapped against the reference using STAR (v2.7.6a and 2.7.7a;^55^) and splice sites information from the gene models. Uniquely mapped reads were converted into counts per gene model and sample using featureCounts (v2.0.1;^56^).

#### Bulk RNA-seq analysis

The iDEP.93 web application (http://bioinformatics.sdstate.edu/idep/)^57^ was used for bullk RNA-seq analysis. Raw counts were filtered for a minimum of 0.1 counts per million in at least one sample, and subsequently normalised using EdgeR, as log_2_(counts per million +1) with missing values treated as 0. Differential gene expression (DGE) analysis was performed using DESeq2 method, with senescent and non-senescent cells paired for individual animals.

#### Gene set enrichment analysis

Gene set enrichment analysis^58^ was performed using the GSEA 4.2.3 software (Broad Institute) using pre-ranked gene lists (given in Table S3) determined from DGE analysis as described above.

#### Single-cell RNA-seq analysis and dataset integrations

Three single cell raw count matrixes were produced: two from internally produced datasets (sorted blastema senescent cells, and blastema macrophages) and one from published dataset (Gerber et. al, blastema and limb total cells,^35^). The blastema macrophage dataset was generated by FACS sorting mCherry+ macrophages from 2 mid-bud blastemas from a *TgTol2(Dr*.*mpeg:mCherry)*^MHY^ transgenic axolotl. Filtering and processing steps were performed on R notebooks with the use of Seurat package functions^59^. In addition, Scanpy functions were used for the calculation of markers and plots by custom Python notebooks^60^. For internal datasets, cells expressing less than 500 genes or less than 9000 UMIs were filtered out. An upper threshold was applied to filter out any clear outliers in the data (more than 20K or 100k expressed genes per cell). Data from Gerber et al. was subsetted to include only connective tissue cells from the blastema and filtered for cells expressing at least 250 genes. Counts in the three datasets were transformed by SCTransform Seurat function and regressed by total UMI content. Dimensionality reduction was obtained by the UMAP method run on the first 30 principal components. Clusters were calculated using the Louvain method with resolution set to 1, on the neighbour network computed with default parameters. Integration of senescent macrophage and connective tissue data with total blastema data were obtained by the anchor-based integration approach implemented in Seurat using 4k features. All the notebooks used for processing raw files and producing the final sc-analysis figures can be found at https://github.com/yun-crtd/Yu-et-al-2022.

#### Ribosomal RNA analysis

We obtained gene lists of significantly upregulated genes in bSCs for early, mid, late and palette stages from bulk RNA-seq datasets. The intersection of these lists comprised 61 commonly upregulated genes conserved across all stages, of which 5/61 corresponded to rRNA transcripts, leading to the identification of ribosomal RNAs as a core signature of senescent cells in vivo. Subsequently, ribosomal regions 18S, 5’ETS, ITS1, and ITS2 were queried against the AmexG_v6.0-DD reference assembly with a local installation of NCBI BLAST 2.13.0 (blastn), parametrized with E-value < 10^−4^ and a word size of 11. Hits were filtered to keep those with at least 90% query coverage. Per sample, the raw read count mapping to each hit, from blastn, were retrieved from corresponding bam files using samtools-bedcov (1.15.1). Then, raw read counts at each coordinate were normalized with EdgeR 3.36.0 on R 4.1.3, as log_2_(counts per million +1) with missing values treated as 0. Visualizations were carried out with ggplot2 3.3.5.

#### qPCR analysis of rRNA processing

Quantitative PCR analysis of rRNA processing was performed using 1X SYBR Green supermix (BioRad) and 500 nM forward and reverse primers, and Rho+ and Rho-cell-derived cDNA as templates. Each sample was run in triplicate. Primers used for qRT-PCR are shown in Key Resources Table - Oligonucleotides.

## QUANTIFICATION AND STATISTICAL ANALYSIS

Animals in each sample group were randomly allocated. Statistical tests employed, sample group size (n), mean, dispersion and precision measures (SD and/or SEM) are indicated in each figure or figure legend. All experiments were carried out in at least three biological replicates. Statistical analyses (e.g. paired or unpaired two-tailed t-test, two-way ANOVA with Sidak’s multiple comparisons) were performed using GraphPad Prism 8 software.

## SUPPLEMENTAL TABLES

**Table S1. DGE analysis of proliferating and senescent AL1 cells (separate Excel file), related to Figure 3**. Results of DGE analysis using DESeq2. Genes are listed in rows. The first 3 columns list gene name, log2-fold change between senescent versus non-senescent cells, and adjusted p values. The following columns provide EdgeR normalised counts for proliferating and senescent AL1 cells.

**Table S2. DGE analysis of non-senescent and senescent blastema cells at early, mid, late and palette stages (separate Excel file), related to Figure 3**. Results of DGE analysis using DESeq2. Genes are listed in rows. The first 3 columns list gene name, log2-fold change between senescent versus non-senescent cells, and adjusted p values. The following columns provide EdgeR normalised counts for proliferating and senescent blastema cells.

**Table S3. Intersection of commonly upregulated genes between *in vitro* and *in vivo* senescent cells and GSEA gene list (separate Excel file), related to Figure 3**. Sheet 1 shows significantly upregulated genes in senescent AL1 cells, endogenous bSCs identified through DESeq2 with an FDR cut-off of 0.1 and a minimum fold change of 1.5, and results from ribosomal BLAST query. Sheet 2 lists commonly upregulated genes shared between senescent AL1s and bSCs, highlighting ribosomal transcripts. Related to Figure 3E. Sheet 3 onwards: lists the genes used for GSEA on senescent and non-senescent cells, and their sources.

**Table S4. Marker gene lists used for cluster annotation of bSC scRNA-seq (separate Excel file), related to Figure 3**. Sheet 1 provides selected marker genes used for annotation of bSC scRNA-seq clusters. Clusters with expression of 45S rRNA were designated as deep senescent cells. Sheet 2 provides full marker gene lists of all clusters. Cluster correspond to those shown in Fig. 3, J-K and Fig. S8A.

## REFERENCES

1. Brockes, J.P., and Kumar, A. (2008). Comparative aspects of animal regeneration. Annual review of cell and developmental biology 24, 525–549.10.1146/annurev.cellbio.24.110707.175336.

2. Cox, B.D., Yun, M.H., and Poss, K.D. (2019). Can laboratory model systems instruct human limb regeneration? Development 146.10.1242/dev.181016.

3. Tanaka, E.M. (2016). The Molecular and Cellular Choreography of Appendage Regeneration. Cell 165, 1598–1608.10.1016/j.cell.2016.05.038.

4. Yun, M.H., Davaapil, H., and Brockes, J.P. (2015). Recurrent turnover of senescent cells during regeneration of a complex structure. eLife 4.10.7554/eLife.05505.

5. Acosta, J.C., O’Loghlen, A., Banito, A., Guijarro, M.V., Augert, A., Raguz, S., Fumagalli, M., Da Costa, M., Brown, C., Popov, N., et al. (2008). Chemokine signaling via the CXCR2 receptor reinforces senescence. Cell 133, 1006–1018.10.1016/j.cell.2008.03.038S0092-8674(08)00619-3 [pii].

6. Campisi, J. (2013). Aging, cellular senescence, and cancer. Annu Rev Physiol 75, 685–705.10.1146/annurev-physiol-030212-183653.

7. Coppe, J.P., Patil, C.K., Rodier, F., Sun, Y., Munoz, D.P., Goldstein, J., Nelson, P.S., Desprez, P.Y., and Campisi, J. (2008). Senescence-associated secretory phenotypes reveal cell-nonautonomous functions of oncogenic RAS and the p53 tumor suppressor. PLoS Biol 6, 2853–2868.10.1371/journal.pbio.0060301.

8. Kuilman, T., Michaloglou, C., Vredeveld, L.C., Douma, S., van Doorn, R., Desmet, C.J., Aarden, L.A., Mooi, W.J., and Peeper, D.S. (2008). Oncogene-induced senescence relayed by an interleukin-dependent inflammatory network. Cell 133, 1019–1031.10.1016/j.cell.2008.03.039.

9. Gorgoulis, V., Adams, P.D., Alimonti, A., Bennett, D.C., Bischof, O., Bishop, C., Campisi, J., Collado, M., Evangelou, K., Ferbeyre, G., et al. (2019). Cellular Senescence: Defining a Path Forward. Cell 179, 813–827.10.1016/j.cell.2019.10.005.

10. Birch, J., and Gil, J. (2020). Senescence and the SASP: many therapeutic avenues. Genes Dev 34, 1565–1576.10.1101/gad.343129.120.

11. Binet, F., Cagnone, G., Crespo-Garcia, S., Hata, M., Neault, M., Dejda, A., Wilson, A.M., Buscarlet, M., Mawambo, G.T., Howard, J.P., et al. (2020). Neutrophil extracellular traps target senescent vasculature for tissue remodeling in retinopathy. Science (New York, N.Y.) 369.10.1126/science.aay5356.

12. Krizhanovsky, V., Yon, M., Dickins, R.A., Hearn, S., Simon, J., Miething, C., Yee, H., Zender, L., and Lowe, S.W. (2008). Senescence of activated stellate cells limits liver fibrosis. Cell 134, 657–667.10.1016/j.cell.2008.06.049S0092-8674(08)00836-2 [pii].

13. Meyer, K., Hodwin, B., Ramanujam, D., Engelhardt, S., and Sarikas, A. (2016). Essential Role for Premature Senescence of Myofibroblasts in Myocardial Fibrosis. J Am Coll Cardiol 67, 2018–2028.10.1016/j.jacc.2016.02.047.

14. Ritschka, B., Storer, M., Mas, A., Heinzmann, F., Ortells, M.C., Morton, J.P., Sansom, O.J., Zender, L., and Keyes, W.M. (2017). The senescence-associated secretory phenotype induces cellular plasticity and tissue regeneration. Genes Dev 31, 172–183.10.1101/gad.290635.116.

15. Sarig, R., Rimmer, R., Bassat, E., Zhang, L., Umansky, K.B., Lendengolts, D., Perlmoter, G., Yaniv, K., and Tzahor, E. (2019). Transient p53-Mediated Regenerative Senescence in the Injured Heart. Circulation 139, 2491–2494.10.1161/CIRCULATIONAHA.119.040125.

16. Demaria, M., Ohtani, N., Youssef, S.A., Rodier, F., Toussaint, W., Mitchell, J.R., Laberge, R.M., Vijg, J., Van Steeg, H., Dolle, M.E., et al. (2014). An essential role for senescent cells in optimal wound healing through secretion of PDGF-AA. Dev Cell 31, 722–733.10.1016/j.devcel.2014.11.012S1534-5807(14)00729-1 [pii].

17. Jun, J.I., and Lau, L.F. (2010). The matricellular protein CCN1 induces fibroblast senescence and restricts fibrosis in cutaneous wound healing. Nat Cell Biol 12, 676–685.10.1038/ncb2070.

18. Davaapil, H., Brockes, J.P., and Yun, M.H. (2017). Conserved and novel functions of programmed cellular senescence during vertebrate development. Development 144, 106–114.10.1242/dev.138222.

19. Munoz-Espin, D., Canamero, M., Maraver, A., Gomez-Lopez, G., Contreras, J., Murillo-Cuesta, S., Rodriguez-Baeza, A., Varela-Nieto, I., Ruberte, J., Collado, M., and Serrano, M. (2013). Programmed cell senescence during mammalian embryonic development. Cell 155, 1104–1118.10.1016/j.cell.2013.10.019S0092-8674(13)01295-6 [pii].

20. Storer, M., Mas, A., Robert-Moreno, A., Pecoraro, M., Ortells, M.C., Di Giacomo, V., Yosef, R., Pilpel, N., Krizhanovsky, V., Sharpe, J., and Keyes, W.M. (2013). Senescence is a developmental mechanism that contributes to embryonic growth and patterning. Cell 155, 1119–1130.10.1016/j.cell.2013.10.041S0092-8674(13)01359-7 [pii].

21. Villiard, E., Denis, J.F., Hashemi, F.S., Igelmann, S., Ferbeyre, G., and Roy, S. (2017). Senescence gives insights into the morphogenetic evolution of anamniotes. Biol Open 6, 891–896.10.1242/bio.025809.

22. Rodier, F., and Campisi, J. (2011). Four faces of cellular senescence. J Cell Biol 192, 547–556.10.1083/jcb.201009094.

23. Muñoz-Espín, D., and Serrano, M. (2014). Cellular senescence: from physiology to pathology. Nature reviews. Molecular cell biology 15, 482–496.10.1038/nrm3823.

24. Yun, M.H. (2018). Cellular senescence in tissue repair: every cloud has a silver lining. The International journal of developmental biology 62, 591–604.10.1387/ijdb.180081my.

25. Da Silva-Álvarez, S., Guerra-Varela, J., Sobrido-Cameán, D., Quelle, A., Barreiro-Iglesias, A., Sánchez, L., and Collado, M. (2020). Cell senescence contributes to tissue regeneration in zebrafish. Aging cell 19, e13052.10.1111/acel.13052.

26. Roy, S., Gardiner, D.M., and Bryant, S.V. (2000). Vaccinia as a tool for functional analysis in regenerating limbs: ectopic expression of Shh. Developmental biology 218, 199–205.10.1006/dbio.1999.9556.

27. Yu, Q., Walters, H.E., and Yun, M.H. (2023). Induction and Characterization of Cellular Senescence in Salamanders. Methods in molecular biology (Clifton, N.J.) 2562, 135–154.10.1007/978-1-0716-2659-7_8.

28. Stocum, D.L. (2017). Mechanisms of urodele limb regeneration. Regeneration (Oxford, England) 4, 159–200.10.1002/reg2.92.

29. Muñoz-Espín, D., Rovira, M., Galiana, I., Giménez, C., Lozano-Torres, B., Paez-Ribes, M., Llanos, S., Chaib, S., Muñoz-Martín, M., Ucero, A.C., et al. (2018). A versatile drug delivery system targeting senescent cells. EMBO molecular medicine 10.10.15252/emmm.201809355.

30. Henras, A.K., Plisson-Chastang, C., O’Donohue, M.F., Chakraborty, A., and Gleizes, P.E. (2015). An overview of pre-ribosomal RNA processing in eukaryotes. Wiley interdisciplinary reviews. RNA 6, 225–242.10.1002/wrna.1269.

31. Lessard, F., Igelmann, S., Trahan, C., Huot, G., Saint-Germain, E., Mignacca, L., Del Toro, N., Lopes-Paciencia, S., Le Calvé, B., Montero, M., et al. (2018). Senescence-associated ribosome biogenesis defects contributes to cell cycle arrest through the Rb pathway. Nat Cell Biol 20, 789–799.10.1038/s41556-018-0127-y.

32. Nishimura, K., Kumazawa, T., Kuroda, T., Katagiri, N., Tsuchiya, M., Goto, N., Furumai, R., Murayama, A., Yanagisawa, J., and Kimura, K. (2015). Perturbation of ribosome biogenesis drives cells into senescence through 5S RNP-mediated p53 activation. Cell reports 10, 1310–1323.10.1016/j.celrep.2015.01.055.

33. Pantazi, A., Quintanilla, A., Hari, P., Tarrats, N., Parasyraki, E., Dix, F.L., Patel, J., Chandra, T., Acosta, J.C., and Finch, A.J. (2019). Inhibition of the 60S ribosome biogenesis GTPase LSG1 causes endoplasmic reticular disruption and cellular senescence. Aging cell 18, e12981.10.1111/acel.12981.

34. Kim, T.H., Leslie, P., and Zhang, Y. (2014). Ribosomal proteins as unrevealed caretakers for cellular stress and genomic instability. Oncotarget 5, 860–871.10.18632/oncotarget.1784.

35. Gerber, T., Murawala, P., Knapp, D., Masselink, W., Schuez, M., Hermann, S., Gac-Santel, M., Nowoshilow, S., Kageyama, J., Khattak, S., et al. (2018). Single-cell analysis uncovers convergence of cell identities during axolotl limb regeneration. Science (New York, N.Y.) 362.10.1126/science.aaq0681.

36. Clevers, H., Loh, K.M., and Nusse, R. (2014). Stem cell signaling. An integral program for tissue renewal and regeneration: Wnt signaling and stem cell control. Science (New York, N.Y.) 346, 1248012.10.1126/science.1248012.

37. Milanovic, M., Fan, D.N.Y., Belenki, D., Däbritz, J.H.M., Zhao, Z., Yu, Y., Dörr, J.R., Dimitrova, L., Lenze, D., Monteiro Barbosa, I.A., et al. (2018). Senescence-associated reprogramming promotes cancer stemness. Nature 553, 96–100.10.1038/nature25167.

38. Alexandre, C., Baena-Lopez, A., and Vincent, J.P. (2014). Patterning and growth control by membrane-tethered Wingless. Nature 505, 180–185.10.1038/nature12879.

39. Cherry, C., Andorko, J.I., Krishnan, K., Mejías, J.C., Nguyen, H.H., Stivers, K.B., Gray-Gaillard, E.F., Ruta, A., Han, J., Hamada, N., et al. (2023). Transfer learning in a biomaterial fibrosis model identifies in vivo senescence heterogeneity and contributions to vascularization and matrix production across species and diverse pathologies. GeroScience.10.1007/s11357-023-00785-7.

40. Moiseeva, V., Cisneros, A., Sica, V., Deryagin, O., Lai, Y., Jung, S., Andrés, E., An, J., Segalés, J., Ortet, L., et al. (2023). Senescence atlas reveals an aged-like inflamed niche that blunts muscle regeneration. Nature 613, 169–178.10.1038/s41586-022-05535-x.

41. Hall, B.M., Balan, V., Gleiberman, A.S., Strom, E., Krasnov, P., Virtuoso, L.P., Rydkina, E., Vujcic, S., Balan, K., Gitlin, I., et al. (2016). Aging of mice is associated with p16(Ink4a)- and beta-galactosidase-positive macrophage accumulation that can be induced in young mice by senescent cells. Aging (Albany NY) 8, 1294–1315.10.18632/aging.100991.

42. Liu, J.Y., Souroullas, G.P., Diekman, B.O., Krishnamurthy, J., Hall, B.M., Sorrentino, J.A., Parker, J.S., Sessions, G.A., Gudkov, A.V., and Sharpless, N.E. (2019). Cells exhibiting strong p16(INK4a) promoter activation in vivo display features of senescence. Proceedings of the National Academy of Sciences of the United States of America 116, 2603–2611.10.1073/pnas.1818313116.

43. Sun, Y., Campisi, J., Higano, C., Beer, T.M., Porter, P., Coleman, I., True, L., and Nelson, P.S. (2012). Treatment-induced damage to the tumor microenvironment promotes prostate cancer therapy resistance through WNT16B. Nature medicine 18, 1359–1368.10.1038/nm.2890.

44. Pérez-Garijo, A., Shlevkov, E., and Morata, G. (2009). The role of Dpp and Wg in compensatory proliferation and in the formation of hyperplastic overgrowths caused by apoptotic cells in the Drosophila wing disc. Development 136, 1169–1177.10.1242/dev.034017.

45. Glotzer, G.L., Tardivo, P., and Tanaka, E.M. (2022). Canonical Wnt signaling and the regulation of divergent mesenchymal Fgf8 expression in axolotl limb development and regeneration. eLife 11.10.7554/eLife.79762.

46. Lovely, A.M., Duerr, T.J., Qiu, Q., Galvan, S., Voss, S.R., and Monaghan, J.R. (2022). Wnt Signaling Coordinates the Expression of Limb Patterning Genes During Axolotl Forelimb Development and Regeneration. Frontiers in cell and developmental biology 10, 814250.10.3389/fcell.2022.814250.

47. Nacu, E., Gromberg, E., Oliveira, C.R., Drechsel, D., and Tanaka, E.M. (2016). FGF8 and SHH substitute for anterior-posterior tissue interactions to induce limb regeneration. Nature 533, 407–410.10.1038/nature17972.

48. Lehmann, J., Narcisi, R., Franceschini, N., Chatzivasileiou, D., Boer, C.G., Koevoet, W., Putavet, D., Drabek, D., van Haperen, R., de Keizer, P.L.J., et al. (2022). WNT/beta-catenin signalling interrupts a senescence-induction cascade in human mesenchymal stem cells that restricts their expansion. Cellular and molecular life sciences : CMLS 79, 82.10.1007/s00018-021-04035-x.

49. Bird, T.G., Müller, M., Boulter, L., Vincent, D.F., Ridgway, R.A., Lopez-Guadamillas, E., Lu, W.Y., Jamieson, T., Govaere, O., Campbell, A.D., et al. (2018). TGFβ inhibition restores a regenerative response in acute liver injury by suppressing paracrine senescence. Science translational medicine 10.10.1126/scitranslmed.aan1230.

50. Ferreira-Gonzalez, S., Lu, W.Y., Raven, A., Dwyer, B., Man, T.Y., O’Duibhir, E., Lewis, P.J.S., Campana, L., Kendall, T.J., Bird, T.G., et al. (2018). Paracrine cellular senescence exacerbates biliary injury and impairs regeneration. Nat Commun 9, 1020.10.1038/s41467-018-03299-5.

51. Hoare, M., Ito, Y., Kang, T.W., Weekes, M.P., Matheson, N.J., Patten, D.A., Shetty, S., Parry, A.J., Menon, S., Salama, R., et al. (2016). NOTCH1 mediates a switch between two distinct secretomes during senescence. Nat Cell Biol 18, 979–992.10.1038/ncb3397.

52. Teo, Y.V., Rattanavirotkul, N., Olova, N., Salzano, A., Quintanilla, A., Tarrats, N., Kiourtis, C., Müller, M., Green, A.R., Adams, P.D., et al. (2019). Notch Signaling Mediates Secondary Senescence. Cell reports 27, 997-1007.e1005.10.1016/j.celrep.2019.03.104.

53. Oliveira, C.R., Knapp, D., Elewa, A., Gerber, T., Gonzalez Malagon, S.G., Gates, P.B., Walters, H.E., Petzold, A., Arce, H., Cordoba, R.C., et al. (2022). Tig1 regulates proximo-distal identity during salamander limb regeneration. Nature Communications 13, 1141.10.1038/s41467-022-28755-1.

54. Schindelin, J., Arganda-Carreras, I., Frise, E., Kaynig, V., Longair, M., Pietzsch, T., Preibisch, S., Rueden, C., Saalfeld, S., Schmid, B., et al. (2012). Fiji: an open-source platform for biological-image analysis. Nature methods 9, 676–682.10.1038/nmeth.2019.

55. Dobin, A., and Gingeras, T.R. (2016). Optimizing RNA-Seq Mapping with STAR. Methods in molecular biology (Clifton, N.J.) 1415, 245–262.10.1007/978-1-4939-3572-7_13.

56. Liao, Y., Smyth, G.K., and Shi, W. (2019). The R package Rsubread is easier, faster, cheaper and better for alignment and quantification of RNA sequencing reads. Nucleic acids research 47, e47.10.1093/nar/gkz114.

57. Ge, S.X., Son, E.W., and Yao, R. (2018). iDEP: an integrated web application for differential expression and pathway analysis of RNA-Seq data. BMC bioinformatics 19, 534.10.1186/s12859-018-2486-6.

58. Subramanian, A., Tamayo, P., Mootha, V.K., Mukherjee, S., Ebert, B.L., Gillette, M.A., Paulovich, A., Pomeroy, S.L., Golub, T.R., Lander, E.S., and Mesirov, J.P. (2005). Gene set enrichment analysis: a knowledge-based approach for interpreting genome-wide expression profiles. Proceedings of the National Academy of Sciences of the United States of America 102, 15545–15550.10.1073/pnas.0506580102.

59. Hao, Y., Hao, S., Andersen-Nissen, E., Mauck, W.M., 3rd, Zheng, S., Butler, A., Lee, M.J., Wilk, A.J., Darby, C., Zager, M., et al. (2021). Integrated analysis of multimodal single-cell data. Cell 184, 3573-3587.e3529.10.1016/j.cell.2021.04.048.

60. Wolf, F.A., Angerer, P., and Theis, F.J. (2018). SCANPY: large-scale single-cell gene expression data analysis. Genome biology 19, 15.10.1186/s13059-017-1382-0.

