## Supplemental material combined for "Cellular senescence promotes progenitor cell expansion during axolotl limb regeneration"

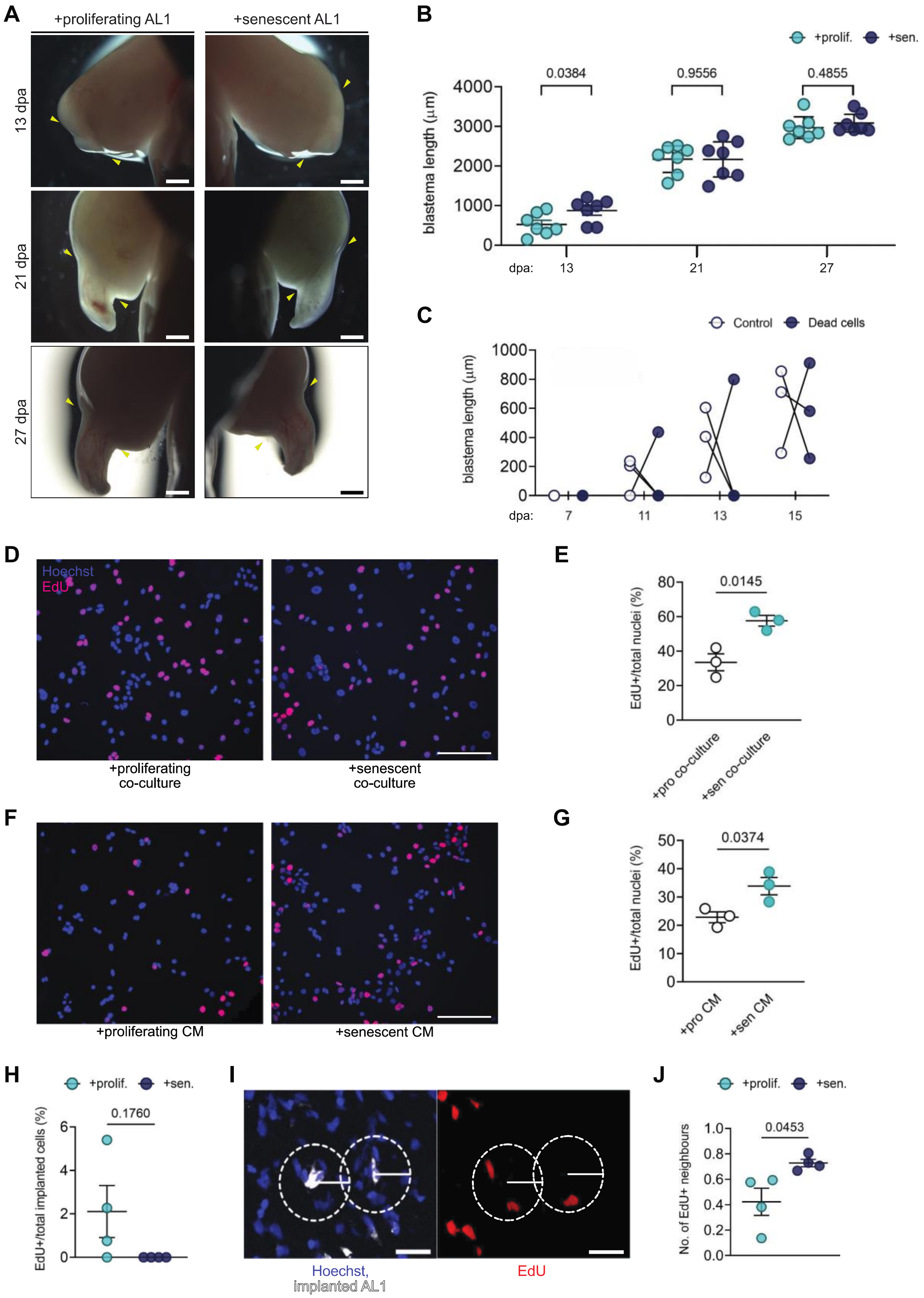
A
+proliferating AL1
+senescent AL1
13 dpa
21 dpa
27 dpa
B
+prolif.
+sen.
blastema length (µm)
0.0384
0.9556
0.4855
dpa: 13 21 27
C
Control
Dead cells
blastema length (µm)
dpa: 7 11 13 15
D
Hoechst
EdU
+proliferating co-culture
+senescent co-culture
E
EdU+/total nuclei (%)
0.0145
+pro co-culture
+sen co-culture
F
+proliferating CM
+senescent CM
G
EdU+/total nuclei (%)
0.0374
+pro CM
+sen CM
H
+prolif.
+sen.
EdU+/total implanted cells (%)
0.1760
I
Hoechst, implanted AL1
EdU
J
+prolif.
+sen.
No. of EdU+ neighbours
0.0453

**Supplementary Fig. 1 (Related to Figure 1). Senescent AL1 cells enhance limb regeneration by stimulating proliferation in a paracrine manner.**

**(A-B)** Dynamics of regeneration during AL1 cell implantation. **(A)** Representative bright-field images of regenerating limbs at the indicated times after implantation of proliferating (left panels) or senescent (right panels) AL1 cells. Scale bar = 1000 μm. **(B)** Quantification of blastema length. *P* values determined by paired two-tailed t-test. Error bars depict mean ± SEM.

**(C)** Implantation of dead cells does not affect the rate of limb regeneration. Quantification of blastema length from regenerating limbs implanted with either control AL1 or ethanol-fixed AL1 cells at indicated days post amputation (dpa). No significant differences observed in blastema length at any timepoint as determined by paired two-tailed t-tests.

**(D-E)** Co-culture with senescent cells promotes proliferation *in vitro*. **(D)** Representative AL1 cultures co-cultured either with proliferating- or senescent-AL1 cells and stained for EdU. Scale bar = 300 μm Blue: Hoechst-stained nuclei; magenta: EdU. **(E)** Quantification of EdU+/total nuclei reveals significantly higher levels of EdU-incorporation in cells co-cultured with senescent AL1 cells. Each datapoint represents a technical replicate. *P* values determined by paired two-tailed t-test. Error bars depict mean ± SEM.

**(F-G)** Senescent cell conditioned media promotes proliferation *in vitro*. **(F)** Representative AL1 cultures treated with either proliferating- or senescent-derived conditioned media and stained for EdU. Scale bar = 300 μm. Blue: Hoechst-stained nuclei; magenta: EdU. CM = conditioned media. **(G)** Quantification of EdU+/total nuclei reveals significantly higher levels of EdU-incorporation in cells cultured with senescent-derived conditioned media. Each datapoint represents a technical replicate. *P* values determined by paired two-tailed t-test. Error bars depict mean ± SEM.

**(H)** Quantification of proliferation of implanted cells. Implanted cells were identified through histology using Vybrant CM labelling DiI and the percentage of EdU+/total implanted cells quantified per blastema. No EdU+ nuclei were detected in implanted senescent AL1 cells.

**(I-J)** Implanted senescent AL1 cells promote proliferation of blastema progenitors in a short-ranged manner. **(I)** Histology of 16 dpa blastemas implanted with AL1 cells. Implanted cells were identified by Vybrant CM DiI labelling and tissue collected at 16 dpa. Average number of EdU+ cells within a 50 μm radius was assessed. **(J)** Quantification of number EdU+ neighbouring cells, based on **(I)**. *P* values determined by paired two-tailed t-test. Error bars depict mean ± SEM.

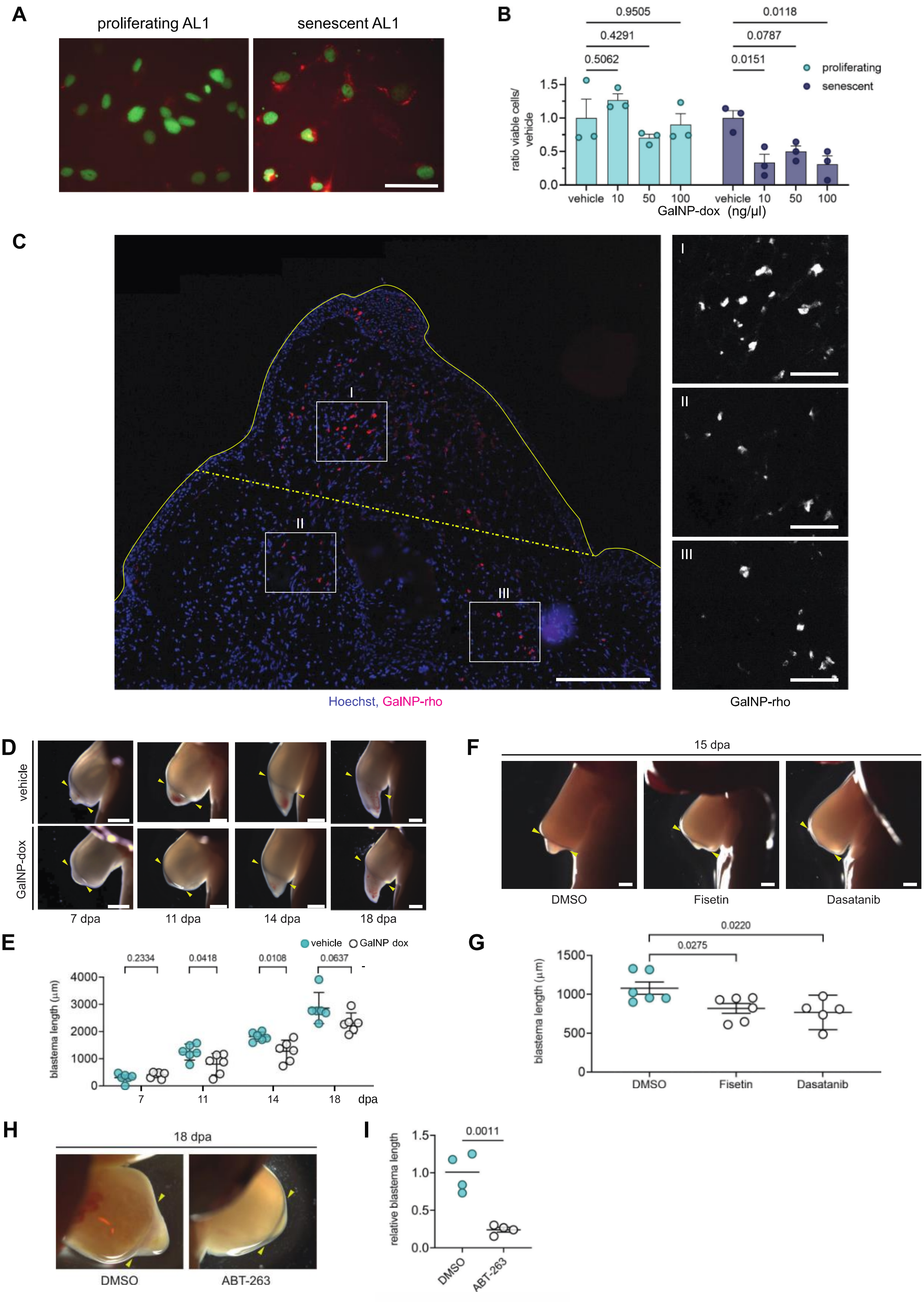
A
proliferating AL1
senescent AL1
B
0.9505
0.4291
0.5062
0.0118
0.0787
0.0151
proliferating
senescent
ratio viable cells/ vehicle
1.5
1.0
0.5
0.0
vehicle 10 50 100 vehicle 10 50 100
GalNP-dox (ng/µl)
C
I
II
III
Hoechst, GalNP-rho
GalNP-rho
D
vehicle
GalNP-dox
7 dpa
11 dpa
14 dpa
18 dpa
E
vehicle
GalNP dox
0.2334
0.0418
0.0108
0.0637
blastema length (µm)
4000
3000
2000
1000
0
7
11
14
18
dpa
F
15 dpa
DMSO
Fisetin
Dasatanib
G
0.0220
0.0275
blastema length (µm)
1500
1000
500
0
DMSO
Fisetin
Dasatanib
H
18 dpa
DMSO
ABT-263
I
0.0011
relative blastema length
1.5
1.0
0.5
0.0
DMSO
ABT-263

**Supplementary Fig. 2 (Related to Figure 2). Elimination of senescent cells through senescence-specific GalNP or senolytic treatments attenuates regeneration.**

**(A)** GalNP-rho specifically labels senescent cells *in vitro*. Representative images of proliferating or senescent AL1 cells *in vitro* treated with a 24 h pulse of GalNP-rho. Green: Hoechst; red: rhodamine. Scale bar = 50 μm.

**(B)** GalNP system enables specific killing of senescent cells *in vitro*. Number of live cells as assessed by calcein green AM staining for indicated doses of GalNP-dox in proliferating or senescent AL1 cells *in vitro*. Data displayed as ratio of live cells normalised against vehicle treatment. *P*-values determined by unpaired two-tailed t-test. Data points represent technical replicates.

**(C)** Representative sections of 15 dpa blastemas at 48 h after GalNP-rho injection. Solid yellow line: outline of blastema; yellow dotted line: amputation plane. Blue: Hoechst; red: rhodamine. Scale bar = 500 μm for main; 50 μm for insets.

**(D-E)** Dynamics of regeneration during GalNP-mediated depletion. **(D)** Bright-field images of regenerating limbs at indicated timepoints, treated with vehicle (top panels) or GalNP-dox (bottom panels). Scale bar = 1000 μm. Arrowheads indicate amputation site. **(E)** Quantification of blastema length across indicated times. *P* values determined by unpaired two-tailed t-tests. Error bars depict mean ± SEM.

**(F)** Bright-field images of regenerating limbs at 15 dpa treated with DMSO, or indicated senolytics. Scale bar = 1000 μm.

**(G)** Quantification of blastema length, based on **(C)**. *P* values determined by unpaired two-tailed t-test against DMSO-treated animals. Error bars depict mean ± SEM.

**(H)** Bright-field images of regenerating limbs at 18 dpa treated with DMSO or ABT-263.

**(I)** Quantification of blastema length, based on **(H)**. Data represented as relative blastema length normalised against mean DMSO blastema length. *P* values determined by unpaired two-tailed t-test. Error bars depict mean ± SEM.

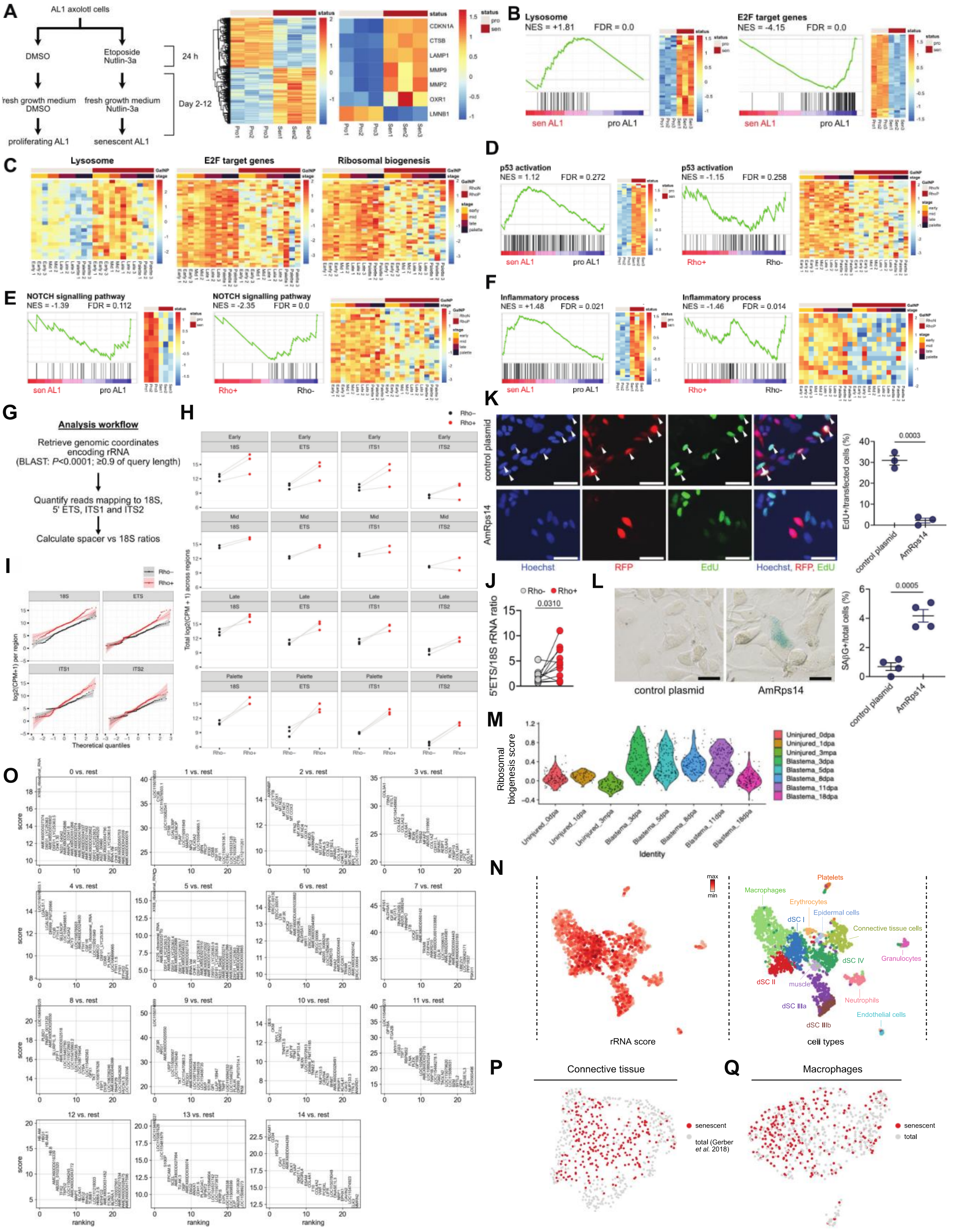

A
AL1 axolotl cells
DMSO
Etoposide Nutlin-3a
24 h
fresh growth medium DMSO
fresh growth medium Nutlin-3a
Day 2-12
proliferating AL1
senescent AL1
status
pro
sen
CDKN1A
CTSB
LAMP1
MMP9
MMP2
OXR1
LMNB1
Pro1
Pro2
Pro3
Sen1
Sen2
Sen3
B
Lysosome
NES = +1.81
FDR = 0.0
sen AL1
pro AL1
E2F target genes
NES = -4.15
FDR = 0.0
C
Lysosome
E2F target genes
Ribosomal biogenesis
GalNP
stage
RhoN
RhoP
early
mid
late
palette
D
p53 activation
NES = 1.12
FDR = 0.272
p53 activation
NES = -1.15
FDR = 0.258
Rho+
Rho-
E
NOTCH signalling pathway
NES = -1.39
FDR = 0.112
NOTCH signalling pathway
NES = -2.35
FDR = 0.0
F
Inflammatory process
NES = +1.48
FDR = 0.021
Inflammatory process
NES = -1.46
FDR = 0.014
G
Analysis workflow
Retrieve genomic coordinates encoding rRNA
(BLAST: P<0.0001; ≥0.9 of query length)
Quantify reads mapping to 18S, 5' ETS, ITS1 and ITS2
Calculate spacer vs 18S ratios
H
Rho−
Rho+
Early
Mid
Late
Palette
18S
ETS
ITS1
ITS2
Total log2(CPM + 1) across regions
I
Rho−
Rho+
log2(CPM+1) per region
Theoretical quantiles
K
control plasmid
AmRps14
Hoechst
RFP
EdU
Hoechst, RFP, EdU
EdU+/transfected cells (%)
0.0003
J
Rho-
Rho+
5'ETS/18S rRNA ratio
0.0310
L
control plasmid
AmRps14
SAβG+/total cells (%)
0.0005
M
Ribosomal biogenesis score
Uninjured_0dpa
Uninjured_1dpa
Uninjured_3mpa
Blastema_3dpa
Blastema_5dpa
Blastema_8dpa
Blastema_11dpa
Blastema_18dpa
Identity
N
max
min
rRNA score
Platelets
Macrophages
Erythrocytes
Epidermal cells
dSC I
Connective tissue cells
dSC IV
Granulocytes
dSC II
muscle
Neutrophils
dSC IIIa
Endothelial cells
dSC IIIb
cell types
O
0 vs. rest
1 vs. rest
2 vs. rest
3 vs. rest
4 vs. rest
5 vs. rest
6 vs. rest
7 vs. rest
8 vs. rest
9 vs. rest
10 vs. rest
11 vs. rest
12 vs. rest
13 vs. rest
14 vs. rest
score
ranking
P
Connective tissue
senescent
total (Gerber et al. 2018)
Q
Macrophages
senescent
total

**Supplementary Fig. 3 (Related to Figure 3). RNA-seq unveils core features of cellular senescence *in vitro* and *in vivo*.**

**(A)** Schematic of senescence induction in AL1 cells *in vitro* and corresponding heatmaps of differentially expressed genes (DEGs) between proliferating and senescent AL1 cells for total DEGs (left) and senescent-related genes (right).

**(B)** Pre-ranked GSEA plots and corresponding heatmaps for leading-edge genes comparing *in vitro* senescent and non-senescent cells for lysosome and E2F target genes in AL1 cells.

**(C)** Pre-ranked GSEA plots and corresponding heatmaps for leading-edge genes comparing *in vivo* senescent and non-senescent blastemal cells for lysosome, E2F target, and ribosomal biogenesis-associated genes, as in **Fig. 3 D & G**.

**(D-F)** Pre-ranked GSEA plots and corresponding heatmaps for leading-edge genes comparing senescent and non-senescent cells *in vitro* (left panels) and *in vivo* (right panels) for: **(D)** p53 transcriptional target genes; **(E)** NOTCH signalling pathway transcriptional targets; and **(F)** inflammatory process.

**(G)** Overview of RNA processing analysis strategy. BLAST was filtered to return results within 90% of query length to return true ribosomal RNA regions.

**(H)** qqPlots reveal enrichment of rRNA regions in bSCs (Rho+) versus their non-senescent (Rho-) counterparts for all regions analysed (18S, 5' ETS, ITS1 and ITS2).

**(I)** Quantification of total reads mapping to indicated rRNA regions comparing senescent (Rho+) and non-senescent (Rho-) populations, stratified by regeneration stages.

**(J)** Quantitative PCR analysis of unprocessed and processed rRNA ratios, using primers specifically targeting 5' ETS and 18S rRNA regions respectively, against cDNA derived from senescent (Rho+) and non-senescent (Rho-) cells.

**(K)** Overexpression of Rps14 induces strong reductions in cellular proliferation. *Left:* AL1 cells lipofected with either pN2-CMV:RFP control plasmids (top row) or pN2-CMV:AmRps14-RFP plasmids (bottom row) at 6 days post-lipofection. Blue: Hoechst; red: RFP; green: EdU. Scale bar = 100 μm for main. *Right:* Quantification of cell-cycle induction between control and AmRps14-overexpressing AL1 cells. Data represented as percentage of EdU-positive of total transfected cells, as identified by RFP signal. Each data point corresponds to one individual transfection. *P* values determined by unpaired two-tailed t-test. Error bars depict mean ± SEM.

**(L)** Overexpression of Rps14 enforces induction of cellular senescence. *Left:* Representative bright-field images of AL1 cells lipofected with either pN2-CMV:RFP control plasmids or pN2-CMV:Rps14 plasmids at 17 days post-lipofection, stained with SAβG. Scale bar = 40 μm. *Right:* Quantification of senescence induction between control and AmRps14-overexpressing AL1 cells. Data represented as percentage of SAβG-positive cells of total cells. Each data point corresponds to one individual transfection. *P* values determined by unpaired two-tailed t-test. Error bars depict mean ± SEM.

**(M)** Ribosomal biogenesis expression during axolotl limb regeneration. Violin plot representation of ribosomal protein score (Table S3) across mature or limb blastema connective tissue single cells[35] at the indicated days post amputation (dpa). Mpa=months post amputation.

**(N)** UMAP of single sorted bSCs showing overlaid rRNA score expression (left) or cluster annotations.

**(O)** Marker gene expression of individual clusters.

**(P-Q)** Integrated scRNA-seq datasets used to generate plots in Fig. 3, L-M. **(P)** Integration of senescent connective tissue (subsetted from bSCs) with total connective tissue[35]. **(Q)** Integration of senescent macrophages (subsetted from bSCs) with total (*mpeg:mCherry*-sorted) macrophages.

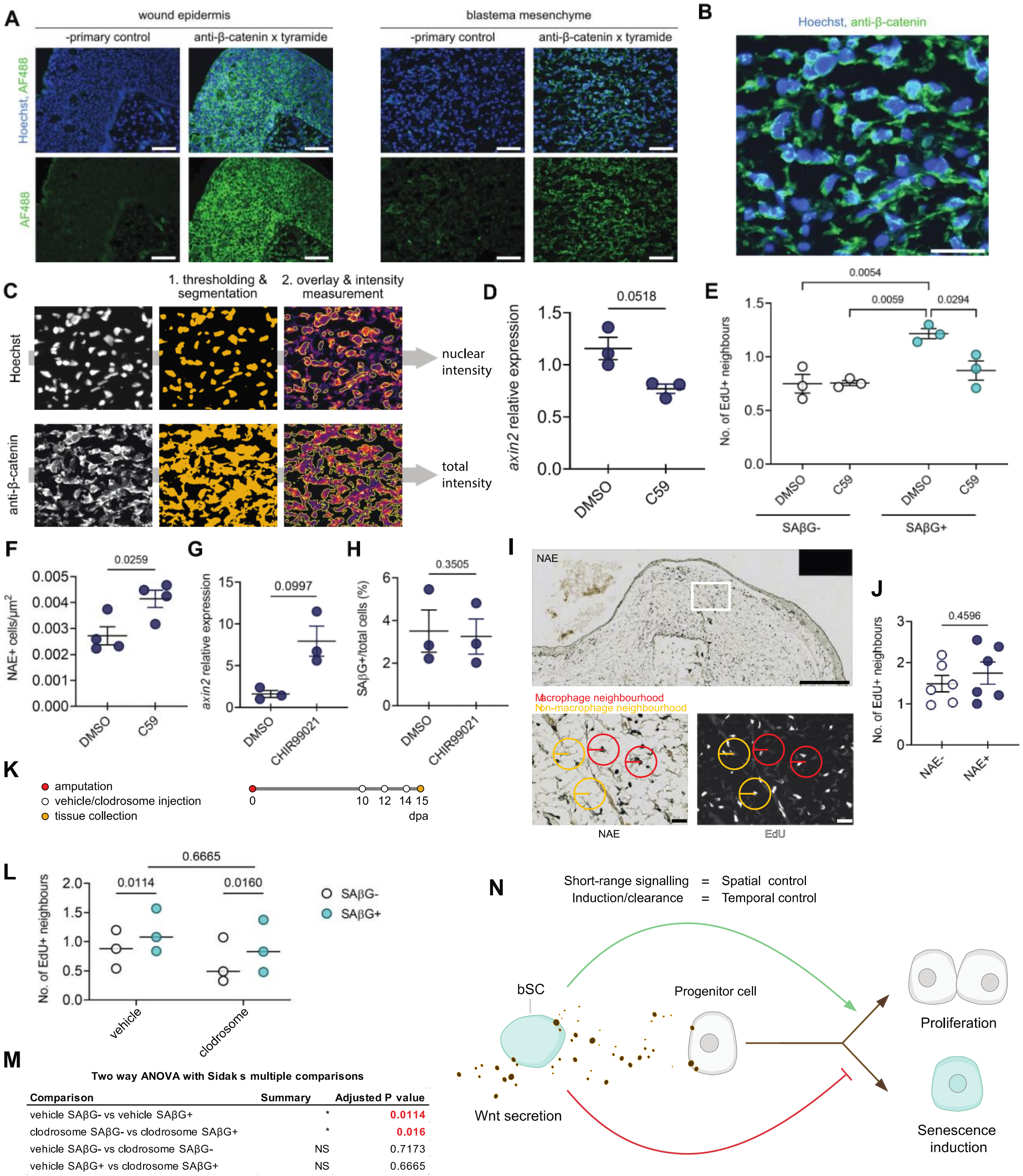


M

**Two way ANOVA with Sidak s multiple comparisons**

| Comparison | Summary | Adjusted P value |
|---|---|---|
| vehicle SAβG- vs vehicle SAβG+ | * | 0.0114 |
| clodrosome SAβG- vs clodrosome SAβG+ | * | 0.016 |
| vehicle SAβG- vs clodrosome SAβG- | NS | 0.7173 |
| vehicle SAβG+ vs clodrosome SAβG+ | NS | 0.6665 |

**Supplementary Fig. 7 (Related to Figure 4). Blastemal senescent cell-derived effects are mediated by Wnt signalling.**

**(A-C)** Analysis of β-catenin nuclear translocation in the axolotl blastema. **(A)** Validation of anti-β-catenin antibody in axolotl tissues. Scale bars = 100 μm. Blue: Hoechst-stained nuclei; green: anti-β-catenin antibody. **(B)** Example input image for quantification used in **(C)**. Scale bar = 50 μm. Blue: Hoechst-stained nuclei; green: anti-β-catenin antibody. **(C)** Left panels: from Fig. S9B, images separated by channels as indicated. Middle panels: result of thresholding and segmentation. Right panels: segmentation results overlaid onto original β-catenin images; β-catenin signal is measured within segmentation-defined boundaries (shown as yellow outline).

**(D)** Validation of C59 efficacy *in vivo*. mRNA levels of *axin2* in DMSO or C59-treated blastemas measured via qRT-PCR. *axin2* expression values calculated using the cycle threshold method (ΔΔCT) method normalised against large ribosomal protein 4 (*Rpl4*). Data points correspond to biological replicates. *P* values determined by unpaired two-tailed t-test. Error bars depict mean ± SEM.

**(E)** Quantification of neighbouring cell proliferation reveals bSC effects are dependent on Wnt signalling. Blastema sections from DMSO or C59-treated animals were co-stained for SAβG and EdU. The average numbers of $EdU^+$ cells surrounding senescent versus non-senescent cells within a 50 μm radius were quantified per section. Data points correspond to biological replicates. *P* values determined by paired two-tailed t-test. Error bars depict mean ± SEM (n=3).

**(F)** Quantification of macrophage recruitment (detected through α-napthtyl esterase [NAE] enzymatic activity) to the blastema mesenchyme following Wnt inhibition using C59. Data is represented as the number of macrophages per $um^2$ of blastema tissue. Data points correspond to biological replicates. *P* values determined by unpaired two-tailed t-test. Error bars depict mean ± SEM (n=4).

**(G)** Validation of CHIR99021 efficacy *in vivo*. mRNA levels of *axin2* in DMSO or CHIR99021-treated blastemas derived from contralateral limbs measured via qRT-PCR. *axin2* expression values calculated using the cycle threshold method (ΔΔCT) method normalised against large ribosomal protein 4 (*Rpl4*). Data points correspond to biological replicates. *P* values determined by paired two-tailed t-test. Error bars depict mean ± SEM.

**(H)** Quantification of percentage of SAβG+/total cells within blastema sections in DMSO or CHIR99021-treated blastemas derived from contralateral limbs. *P* values determined by paired two-tailed t-test. Error bars depict mean ± SEM (n=5).

**(I-J)** Macrophages do not exhibit a neighbouring-cell effect. **(I)** Representative sections of 15 dpa blastemas co-stained for EdU and macrophages detected by NAE enzymatic activity. Top panel: bright-field image of overview of blastema. Scale bar = 500 μm. Bottom panels: close ups of region demarcated by white box; bottom left – bright-field; bottom right – EdU; scale bar = 50 μm. **(J)** Quantification of neighbouring cell proliferation reveals macrophages do not exhibit a neighbouring cell effect. 15 dpa blastema sections were co-stained for NAE and EdU. The average numbers of $EdU^+$ cells surrounding macrophages versus non-macrophage cells were quantified within a 50 μm radius. Data points correspond to biological replicates. *P* values determined by paired two-tailed t-test. Error bars depict mean ± SEM (n=6 biological replicates).

**(K-L)** Senescent-cell associated neighbouring cell effect persists after macrophage depletion. **(K)** Experimental timeline for macrophage-depletion using clodrosome treatments. **(L)** Quantification of senescent cell-neighbouring cell proliferation. Blastema sections from vehicle or clodrosome-treated animals were co-stained for SAβG and EdU. The average numbers of $EdU^+$ cells surrounding senescent versus non-senescent cells within a 50 μm radius were quantified per section. Data points correspond to biological replicates. *P* values determined by two-way ANOVA with Sidak's multiple comparisons. Error bars depict mean ± SEM (n=3). **(M)** Table of results of two-way ANOVA with Sidak's multiple comparisons, corresponding to data in **(L)**.

**(N)** A model for the role and regulation of cellular senescence in axolotl limb regeneration. Senescent-derived Wnt promotes the proliferation of progenitor cells in a paracrine manner, whilst simultaneously preventing their induction into the senescence state.

**Table S1. DGE analysis of proliferating and senescent AL1 cells (separate Excel file), related to Figure 3.**
Results of DGE analysis using DESeq2. Genes are listed in rows. The first 3 columns list gene name, log2-fold change between senescent versus non-senescent cells, and adjusted p values. The following columns provide EdgeR normalised counts for proliferating and senescent AL1 cells.

**Table S2. DGE analysis of non-senescent and senescent blastema cells at early, mid, late and palette stages (separate Excel file), related to Figure 3.**
Results of DGE analysis using DESeq2. Genes are listed in rows. The first 3 columns list gene name, log2-fold change between senescent versus non-senescent cells, and adjusted p values. The following columns provide EdgeR normalised counts for proliferating and senescent blastema cells.

**Table S3. Intersection of commonly upregulated genes between *in vitro* and *in vivo* senescent cells and GSEA gene list (separate Excel file), related to Figure 3.**
Sheet 1 shows significantly upregulated genes in senescent AL1 cells, endogenous bSCs identified through DESeq2 with an FDR cut-off of 0.1 and a minimum fold change of 1.5, and results from ribosomal BLAST query. Sheet 2 lists commonly upregulated genes shared between senescent AL1s and bSCs, highlighting ribosomal transcripts. Related to Figure 3E. Sheet 3 onwards: lists the genes used for GSEA on senescent and non-senescent cells, and their sources.

**Table S4. Marker gene lists used for cluster annotation of bSC scRNA-seq (separate Excel file), related to Figure 3.**
Sheet 1 provides selected marker genes used for annotation of bSC scRNA-seq clusters. Clusters with expression of 45S rRNA were designated as deep senescent cells. Sheet 2 provides full marker gene lists of all clusters. Cluster correspond to those shown in Fig. 3, J-K and Fig. S8A.
